# Developmental origins of cell heterogeneity in the human lung

**DOI:** 10.1101/2022.01.11.475631

**Authors:** Alexandros Sountoulidis, Sergio Marco Salas, Emelie Braun, Christophe Avenel, Joseph Bergenstråhle, Marco Vicari, Paulo Czarnewski, Jonas Theelke, Andreas Liontos, Xesus Abalo, Žaneta Andrusivová, Michaela Asp, Xiaofei Li, Lijuan Hu, Sanem Sariyar, Anna Martinez Casals, Burcu Ayoglu, Alexandra Firsova, Jakob Michaëlsson, Emma Lundberg, Carolina Wählby, Erik Sundström, Sten Linnarsson, Joakim Lundeberg, Mats Nilsson, Christos Samakovlis

## Abstract

The lung contains numerous specialized cell-types with distinct roles in tissue function and integrity. To clarify the origins and mechanisms generating cell heterogeneity, we created a first comprehensive topographic atlas of early human lung development. We report 83 cell states, several spatially-resolved developmental trajectories and predict cell interactions within defined tissue niches. We integrated scRNA-Seq and spatial transcriptomics into a web-based, open platform for interactive exploration. To illustrate the utility of our approach we show distinct states of secretory and neuroendocrine cells, largely overlapping with the programs activated either during lung fibrosis or small cell lung cancer progression. We define the origin of uncharacterized airway fibroblasts associated with airway smooth muscle in bronchovascular bundles, and describe a trajectory of Schwann cell progenitors to intrinsic parasympathetic neurons controlling bronchoconstriction. Our atlas provides a rich resource for further research and a reference for defining deviations from homeostatic and repair mechanisms leading to pulmonary diseases.

## Introduction

The traditional account of cellular heterogeneity in the lung based on meticulous histology and the expression of few characteristic markers, suggests more than 40 cell-types in the adult human lung (Franks et al., 2008). These can be roughly classified into endothelial cells lining vessels and capillaries, epithelial cells constructing the airway tree and gas-exchange surface (Junqueira et al.) and stromal cells, such as fibroblasts maintaining structural tissue integrity and supporting the functionality of other cells (Desai et al., 2014; Sveiven and Nordgren, 2020). Resident immune cells, like interstitial and alveolar macrophages, are responsible for pathogen recognition and activation of host defenses (Lloyd and Marsland, 2017; Ural et al., 2020). Chemosensory and mechanosensory neurons from the vagal nerve innervate the lung to control basic physiological functions, including breathing frequency (Chang et al., 2015). The lung cell-type repertoire has been further expanded by recent developments in single-cell genomics allowing the interrogation of hundreds of thousand cells from healthy and diseased human lungs (Adams et al., 2020; Okuda et al., 2021; Travaglini et al., 2020; Vieira Braga et al., 2019). To date, 58 distinct cell-types and states can be categorized into the five major classes of cells in the adult healthy lung. This heterogeneity is intriguing, considering that the organ derives from two simple endoderm-derived buds that branch into a mesenchymal pouch to create the tubular and acinar networks of the airways and alveolar gas exchange surfaces (Morrisey and Hogan, 2010). The surrounding mesoderm produces (i) a boundary layer of mesothelial cells, distal lung mesenchyme progenitors (Hein et al., 2021), airway smooth muscle (ASM), vascular smooth muscle and pericytes (VSM) (Kumar et al., 2014) and (ii) endothelial progenitors (Bautch and Caron, 2015). These early endothelial cells proliferate and migrate to produce highly complex pulmonary lymphatic and endothelial vascular networks (Healy et al., 2000) (Greif et al., 2012). Even if hematopoiesis does not occur in the lung, many mesoderm-derived immune cell-types (Cao et al., 2020; Julien et al., 2016; Popescu et al., 2019) colonize the organ creating its resident immune cells (Hashimoto et al., 2013; Ivarsson et al., 2013). In addition to the vagal neurons, ectoderm-derived, neural crest cells migrate extensively into the developing lung buds and form parasympathetic ganglia containing both neurons and glial cells (Burns et al., 2008; Freem et al., 2010). These structures are located close to the proximal airways (rarely below sub-segmental level) and extend neuronal projections towards the ASM (Burns *et al*., 2008; Cho et al., 2019).

Our knowledge of human lung development largely derives from animal experiments and from simplified organoid cultures (Cao *et al*., 2020; Nikolic et al., 2017) underscoring the lack of systematic studies of intact embryonic tissues. In addition, human tissues are general poorly charted with respect to systematic cell-type compositions and spatial relationships, that go beyond positioning of single cell-types based on co-expression of a handful or less protein and/or mRNA markers. Comprehensive cell maps reveal the topologies of each cell-type relatively to all others, and thereby describe niches and tissue-design rules driven by spatial factors and cell interactions. Here, we have focused on the first gestation trimester and applied state-of-art technologies to capture the gene expression profiles of human embryonic lung cells and to map their transitional states in time and space. We first defined seven main cell categories including, mesenchymal, epithelial, endothelial, immune and neuronal cells, in addition to megakaryocytes and erythroblasts/erythrocytes. Higher resolution analysis of each of the major categories suggested 83 cell-identities, corresponding to stable cell-types and transitional differentiation states, with distinct tissue positions for many of them. To better understand cell interactions during organ development, we defined neighborhoods of spatially related cell-identities and used interactome analyses to confirm already known and predict novel communication patterns. We present an on-line platform integrating the scRNA-seq with the spatial analyses to facilitate interactive exploration of our data on whole lung tissue sections at different developmental stages.

## Results

### Overview of cell heterogeneity in the embryonic lung

We dissected lungs from 17 embryos of both sexes, ranging from 5 to 14 weeks post conception (pcw) (Fig. 1 A, viewer) in approximately weekly intervals. Assuming that the left and right lungs are relatively symmetrical, we used material from the one for scRNA-Seq and processed the other for spatial analyses. Similarly, we aimed to analyze consecutive sections of the same tissues using *in situ* mapping approaches to facilitate independent validation of cluster topologies. We first roughly clustered the 163.236 (163K), high-quality cDNA libraries. This initial step facilitated separation into seven main cell-categories: mesoderm-derived (i) mesenchymal cells, (ii) endothelial cells, (iii) immune cells, (iv) megakaryocytes, (v) erythrocytes (RBC) or erythroblasts, (vi) ectoderm-derived neuronal cells and (vii) endoderm-derived epithelial cells (Suppl. Fig. 1). Next, we dived deeper into each of the main cell-types and re-clustered the corresponding cells, aiming to expose additional cell-states that were hidden in the whole dataset analysis. This revealed an unexpectedly high heterogeneity of 83 distinct cell-states (Fig 1 B). To further explore these proposed states and spatially identify them back on the tissue, we mapped gene expression patterns on tissue sections with Spatial Transcriptomics (ST) at 9 different stages (6, 8.5, 10 and 11.5 pcw) (Stahl et al., 2016). Probabilistic analysis of the Visium data (Stereoscope (Andersson et al., 2020)) largely validated the scRNA-Seq results and spatially mapped 75 of the 83 identified cell clusters (Fig 1 C). The missing 8 clusters correspond to the different cell states in parasympathetic ganglia, which were detected as one neuronal cell-state. Apart from defining the cluster locations within tissues, Stereoscope calculates the probability of each cluster to be represented in every ST-spot, allowing to define cluster-pairs located in the same neighborhood. This enables the identification of distinct cellular niches, in defined anatomical positions (Fig. 1 D, STAR methods). We thus defined 4 neighborhoods containing adjacent cell-clusters with higher probability to interact with each other during development. To explore the communication code among cell-states in each neighborhood, we utilized CellChat, based on the expression of ligands, receptors and cofactors in the resident clusters (Jin et al., 2021) and (ii) Nichenet, which also predicts target gene activation in response to cell communication (example in Fig. 1E) (Browaeys et al., 2020). We further generated a set of probes for 177 genes (146 for HybISS and 31 SCRINSHOT) corresponding to cell-state markers and selective signaling components to validate cell communication events by multiplex HybISS (Gyllborg et al., 2020; Ke et al., 2013) (Fig. 1F) and SCRINSHOT (Sountoulidis et al., 2020). To facilitate accessibility and easy data exploration, we constructed an interactive viewer combining (i) scRNA-Seq data analysis with (ii) cell-type distributions on whole lung sections from four developmental stages, (iii) spatial gene expression patterns (experimentally detected and imputed) and (iv) cellular interactions, focusing on distinct tissue neighborhoods (https://tissuumaps.dckube.scilifelab.se/web/private/HDCA/index.html). Below, we present our analysis of mesenchymal, epithelial and neuronal cells-states and their interactions. Immune and endothelial cells and their communications with pericytes are shown in the supplementary note.

**Figure 1.**
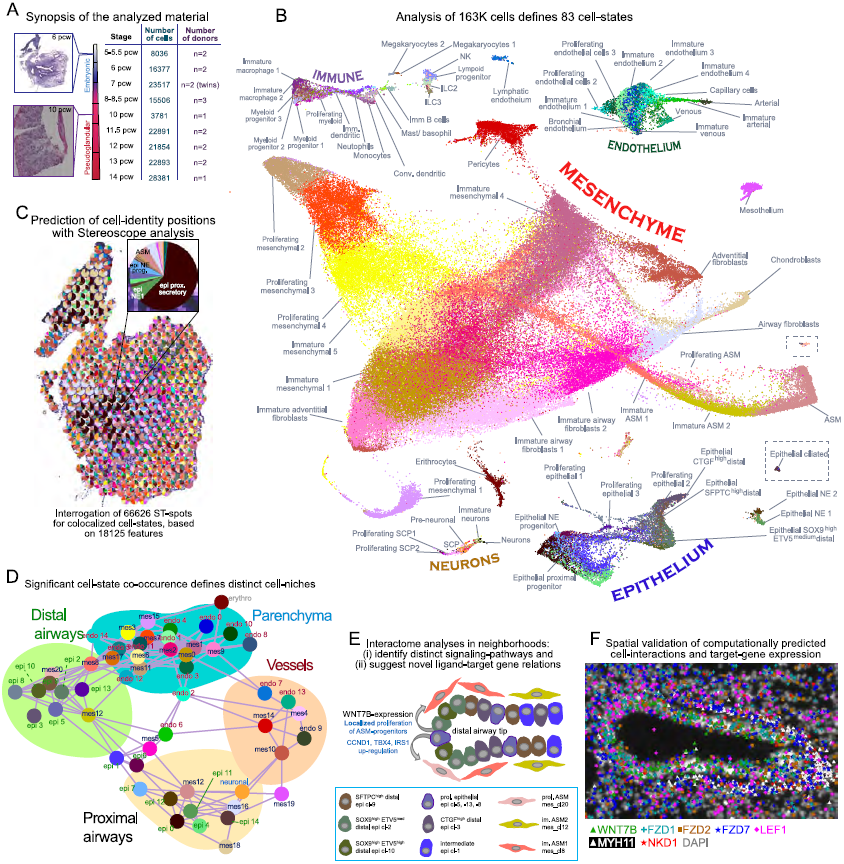
Overview of the study. **(A)** Schematic timeline of the analyzed developmental stages. (Left) Hematoxylin-eosin (H&E) staining images from 6 (top) and 10 (bottom) post-conceptual week (pcw) analyzed lung sections by Spatial Transcriptomics (ST). (Right) Table of analyzed material. First column: developmental stage. Second column: number of analyzed cells after quality controls and third column: number of donors. **(B)** UMAP-plot of the 83 identified cell-clusters by the analyses of the main cell categories (mesenchyme, epithelium, endothelium, immune and neuronal cells). **(C)** Example of an analyzed 6 pcw lung section with ST, showing the Stereoscope cluster positional predictions, with pie-charts (same colors as in “B”). Insert: magnification of an ST-spot, showing its cluster composition. **(D)** Colocalization-graph based on cluster co-occurrence in ST-spots, according to Stereoscope. Neuronal clusters are grouped in a single group (neuronal) and immune cell types are excluded. Line thickness indicates the degree of spatial correlation between two clusters in the ST-spots. Insignificant connections are not shown (Pearson’s *r* <0.04). Distal and proximal airways, vessels and parenchyma are the four identified “cell neighborhoods”. Colors as in “1B”. **(E)** Cartoon of predicted WNT-signaling communication patterns between spatially related clusters (CellChat), showing predictions (NicheNet) of the up-regulated WNT7B-target genes. **(F)** Experimental validation of *WNT7B* communication pattern, between WNT7B^pos^ epithelium and the surrounding mesenchyme, using HybISS.

### Distinct positions for known and novel mesenchymal cell state identities

Mesenchymal cells were the most numerous cell category in our dataset, including ∼138K cells (Suppl. Fig. 1A). Sub-clustering, revealed 6 distinct cell-types and their corresponding immature states, as illustrated by partition-based graph abstraction analysis (PAGA-plot) (Fig. 2A) (Wolf et al., 2019). To annotate the clusters, we used their spatial distribution (examples in Fig. 2B) and the selective expression of known adult mesenchymal cell markers (Suppl. Fig. 2A). We detected: (i) Mesothelial cells at the lung periphery (MS, cl-19) expressing *WT1, MSLN, KRT18* and *KRT19* (Suppl. Fig. 2B), (ii) Pericytes/Vascular Smooth Muscle (VSM, cl-14) associated with endothelial vessels and marked by *PDGFRB, EBF1* and *EGFL6* and moderate levels of *ACTA2* and *TAGLN*, (iii) Chondroblasts (CB, cl-18) expressing *SOX9* and *COL2A1* and surrounding the tracheal and proximal airway epithelium, (iv) Airway Smooth Muscle (ASM, cl-13) expressing *MYH11* and *DACH2* close to airways, (v) adventitial fibroblasts (AdvF, cl-10) expressing *SERPINF1* and *SRFP2* and (vi) airway fibroblasts (AF, cl-16) expressing *ASPN* and *TNC*. AdvF and AF occupied distinct positions in the bronchovascular bundles (Dalpiaz et al., 2017), with the AF cluster being localized closer to airways than AdvF (Fig. 2B). In addition, five of the 21 mesenchymal clusters (cl-3, cl-7, cl-11, cl-16 and cl-20) were composed of proliferating cells and the remaining 10 correspond to immature cell states.

**Figure 2.**
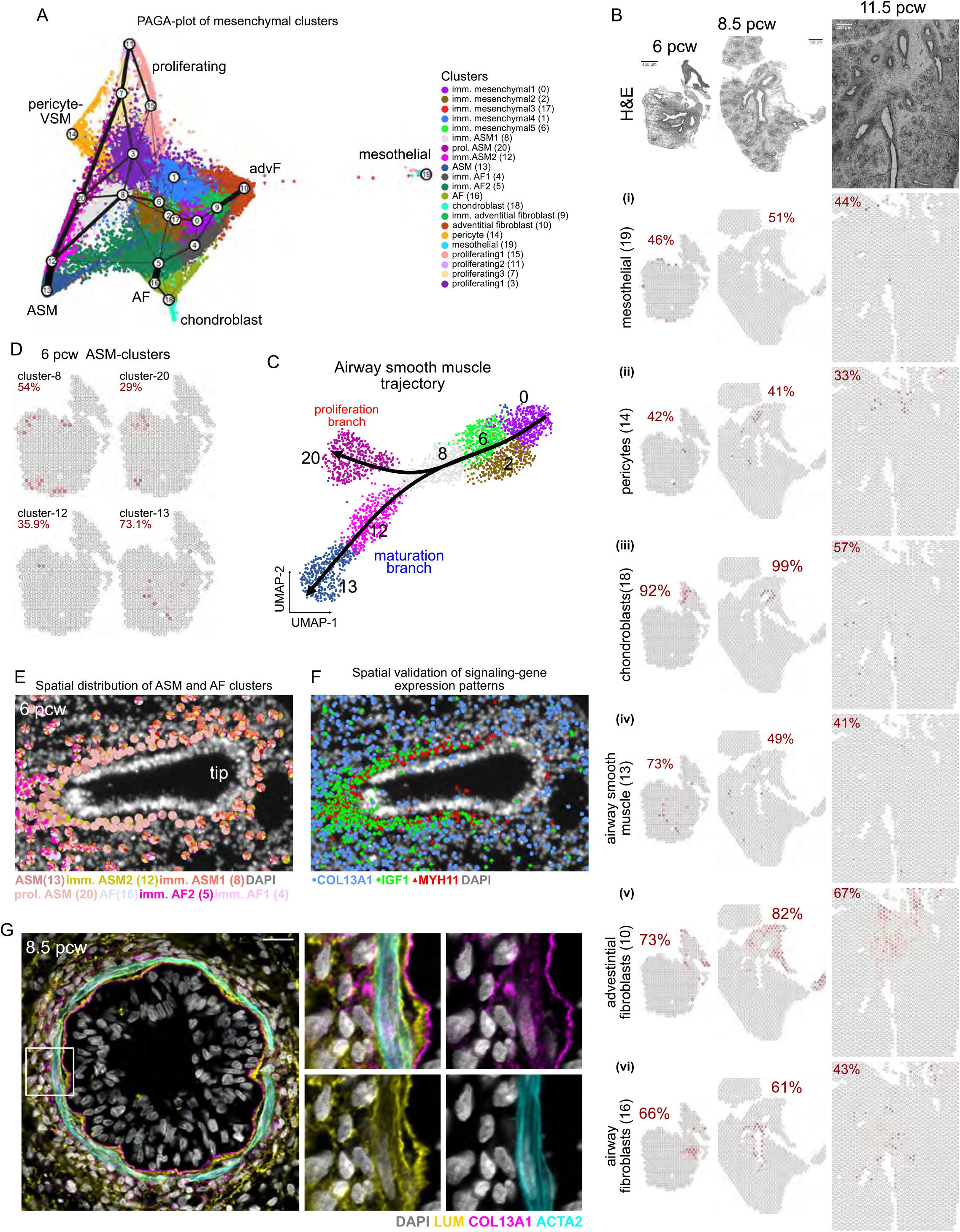
Analysis of mesenchymal cells. **(A)** PAGA-plot of the analyzed mesenchymal cells, superimposed on the UMAP-plot of 138K mesenchymal cells. Line thickness indicates the probability of the cluster connections. Colors show the 21 suggested clusters. **(B)** Stereoscope analysis, based on ST-data, showing the spatial distribution of the developing (i) mesothelial cells (cl-19), (ii) pericytes (cl-14), (iii) chondroblasts (cl-18), (iv) airway smooth muscle (ASM) (cl-13), (v) adventitial fibroblasts (cl-10) and (iii) airway fibroblasts (AF) (cl-16), in 6, 8.5 and 11.5 pcw lung sections. Red numbers correspond to the highest percent value of the indicated cell-type. Dark red: high, gray: zero percent. Tissue structure is shown by H&E staining. Scale-bar: 400 µm. **(C)** Pseudotime analysis of the ASM-trajectory, with Slingshot [3] showing the proliferation (cl-20) and maturation (cl-12 and −13) trajectories. Same colors as in “A”. **(D)** As, in “B” for the ASM-trajectory, in a 6 pcw lung section. **(E)** Spatial localization of the ASM and AF clusters, in a 6 pcw lung section, using probabilistic cell typing (pciSeq) with HybISS data. **(F)** Representative image of *MYH11* (red), *IGF1* (green) and *COL13A1* (blue) detected mRNAs (HybISS) around the same airway, as in “E”. **(G)** Single-plane, confocal-microscopy image of immunofluorescence for COL13A1 (magenta), LUM (yellow) and ACTA2 (cyan), to show AFs and ASM, respectively, in an 8.5 pcw proximal airway (left). Square bracket indicates the area of the images on the right. Nuclear DAPI: gray, Scale-bar 20 µm.

### Airway smooth muscle maturation-states coincide with distinct topologies

The most prominent mesenchymal trajectory, illustrates a differentiation path of immature mesenchymal cells towards ASM. The PAGA-plot connects three immature mesenchymal clusters (cl-0, cl-2 and cl-6) to a proliferating ASM cell cluster (cl-20) and three ASM clusters (−8, −12 and −13) (Fig. 2A). This suggests that the trajectory stems from the cl-0 and through cl-2 and cl-6 connects to the immature ASM-clusters cl-8 and −12, leading to the more mature cl-13. Proliferating ASM cells (mes cl-20) showed high expression of smooth muscle markers, like *ACTA2* and *TAGLN*, implying that they represent a more mature state than mes cl-0 (Suppl. Fig. 2A). Interestingly, cl-20 also selectively expresses extracellular matrix (ECM) proteins (Suppl. Fig. 2C), suggesting that it locally contributes to its composition. We used pseudotime analysis (Street et al., 2018; Van den Berge et al., 2020) to order ASM clusters (Fig. 2C) and define differentially expressed gene-modules that might contribute to ASM differentiation from naïve mesenchymal cells (Suppl. Fig. 2 D-G). Characteristic regulators include the myogenic TF DACH2, which was detected mainly in intermediate states (mes cl-8 and −12) (Suppl. Fig. 2D, 2G: gene module-5). The expression of the NOTCH-ligand, *JAG1* was also increased in these states, in agreement with previous described in vitro analysis (Doi et al., 2006) (Suppl. Fig. 2 F, H). ZEB1 was the most up-regulated transcription factor (TF) in the immature ASM cl-12 (Suppl. Fig. 2G: gene module-6), suggesting that it activates the SM gene program in ASM, as it does in VSM (Nishimura et al., 2006). Differentiation into mature ASM states is proposed to occur in mes cl-12 and cl-13, characterized by increased expression of canonical markers, like *ACTA2, TAGLN* and *MYH11* (Travaglini *et al*., 2020), (Suppl. Fig. 2G: gene module-7). *HHIP*, a target and inhibitor of HH-signaling (Chuang and McMahon, 1999) and the secreted BMP inhibitor GREM2 (Yeung et al., 2014) are enriched in the more mature ASM cluster (Suppl. Fig. 2G: gene modules −7 and −9), implicating the regulation of these pathways in ASM differentiation.

Spatial analysis localized most clusters of this trajectory in distinct positions along the developing airways (Fig 2 D, E), indicating that there is a correlation between the ASM maturation states and topology, with most immature states located peripherally and mature ASM states located closer to proximal airways, similarly to the mouse lung (Kumar *et al*., 2014). Mesenchymal clusters cl-0 and cl-2 were dispersed in the parenchyma (Fig. 1D, Viewer: ST-data) and expressed high *WNT2* and *RSPO2* levels, in contrast to the other immature mesenchymal populations (Suppl. Fig. 2 E, G). This is consistent with defects in ASM differentiation caused by *WNT2* inactivation in mice (Goss et al., 2011). Intriguingly, FGF10, an epithelial outgrowth inducer in mice (Bellusci et al., 1997b), was expressed by cl-0 (Suppl. Fig. 2E) and not by the rest of the ASM-trajectory clusters, which were localized closer to the distal epithelium (Fig. 2D). This suggests that precursors are evenly distributed in the peripheral parenchyma and begin to differentiate close to the bud tip. The analysis of the ASM trajectory suggests that the spatially dispersed *FGF10*^*pos*^ *WNT2*^*pos*^ *RSPO2*^*pos*^ progenitors (mes cl-0 and cl-6) enter the ASM program close to the distal airway buds (mes cl-8) and further proliferate (mes cl-20) and gradually differentiate along the distal to proximal epithelial axis (cl-12 and cl-13) through interactions with epithelial cells and fibroblasts.

### Airway fibroblasts cover the airways, interacting with smooth muscle

Stereoscope analysis showed mes cl-16 to be localized around the airways, as early as 6 pcw (Fig. 2B (vi)). This cluster is negative for *ACTA2* but expresses markers of other adult stromal cell-types, like *ASPN* for myofibroblasts, *SERPINF1* for adventitial fibroblasts (Travaglini *et al*., 2020) and *COL13A1* characterizing a recently described type of lung fibroblast (Raredon et al., 2019) (Suppl. Fig. 2A). Their unique gene expression profile and close proximity to the ASM layer (Fig. 2 E, G) argue that cl-16 corresponds to an undescribed mesenchymal cell-type, which we named “airway fibroblast (AF)”. PAGA -plot analysis suggests a developmental trajectory of these cells involving mesenchymal clusters −4, −5 and leading to cl-16. This is in agreement with the gradual increase of markers like *COL13A1* and *SEMA3E* (Movassagh et al., 2014) in cl-16 (Supp. Fig. 2A). AFs surround the ASMs, with mature AFs (cl-16) being located in proximity to ASM (Fig. 2E-G) and the more immature AF state (cl-5) in more peripheral positions (Fig. 2E, viewer). To explore potential communication routes between them, we focused on signaling pathways emanating from the one and targeting the other (Suppl. Fig. 2I, J). One example is the IGF1-ligand, which is mainly expressed in immature ASM2 (mes cl-12). The corresponding receptor, IGF1R and the predicted target gene *LUM*, are expressed in immature AFs (mes cl-5) (Suppl. Fig. 2K). AFs producing the secreted LUM proteoglycan may thus align and support the formation of collagen bundles around proximal airways (Godoy-Guzman et al., 2012) (Fig. 2G). Additional predicted interactions suggest that WNT5A is produced by ASM cells and induces BMP4 expression through the FZD1 receptor in AFs (Suppl. Fig. 2L). Interestingly, BMP4 is in turn predicted to upregulate *ACTA2* expression in ASM (Wang et al., 2010), suggesting a positive feedback loop, between adjacent AFs and ASMs (Suppl. Fig. 2M). Our results identify AFs as a new cell-type in close contact with ASM cells and suggest their mutual interactions, inducing differentiation and contribution to the organization of connective tissue around the central airways.

### Parasympathetic ganglia and Schwann cell precursor differentiation

The trachea and lungs are innervated by the vagus-nerve containing sympathetic, parasympathetic and sensory neurons. These fibers contain a preganglionic and a postganglionic compartment (De Virgiliis and Di Giovanni, 2020; Netter, 2011). Only parasympathetic ganglia are localized inside the lung, close to the airways, containing the somata of the postganglionic neurons, to innervate ASM (Cho *et al*., 2019) and regulate bronchoconstriction (Netter, 2011). The sources for parasympathetic neurons in mice (Dyachuk et al., 2014; Espinosa-Medina et al., 2014) are neural crest-derived Schwann Cell Precursors (SCPs). *SOX10*^*pos*^ *PHOX2B*^*pos*^ SCPs migrate towards ganglionic positions, where they differentiate to neurons, in a process dependent on the TF ASCL1 (Dyachuk *et al*., 2014).

Sub-clustering of the neuronal cells revealed 8 cell-states, that can be ordered into one main trajectory, corresponding to the transition of Schwann cell progenitors (SCPs) to neurons (Fig. 3 A, B). Clusters −1, −5 and −7 are proliferating SCPs, based on *PCNA* and *MKI67* expression (Bologna-Molina et al., 2013) (Suppl. Fig. 3A). The neuronal clusters cl-0 and cl-3 gradually lose SCP-marker expression while increasing *ASCL1* (Suppl. Fig. 3A), suggesting that they correspond to transient states from SCPs to neurons. cl-2 and cl-6 express the peripheral neuron-specific intermediate filament protein PRPH, *NRG1* and *PHOX2B*. Even if the parasympathetic neuronal markers, nitric oxide synthetase (*NOS1*) and vasoactive intestinal peptide (*VIP*) are minimally expressed, the cl-3, cl-2 and cl-6 cells express both acetylcholine receptors M2 and M3 (*CHRM2* and *CHRM3*) and nicotinic acetylcholine receptor subunits 3 and 7 (*CHRNA3* and *CHRNA7*), in addition to the acetylcholinesterase (*ACHE*) and *SLC5A7*, the latter encoding the high affinity choline transporter for intraneuronal acetylcholine synthesis (Apparsundaram et al., 2000) (Suppl. Fig. 3B). Thus, these cells are parasympathetic neurons, which can respond to acetylcholine but they are still immature, as indicated by NOS1 and VIP minimal expression.

**Figure 3.**
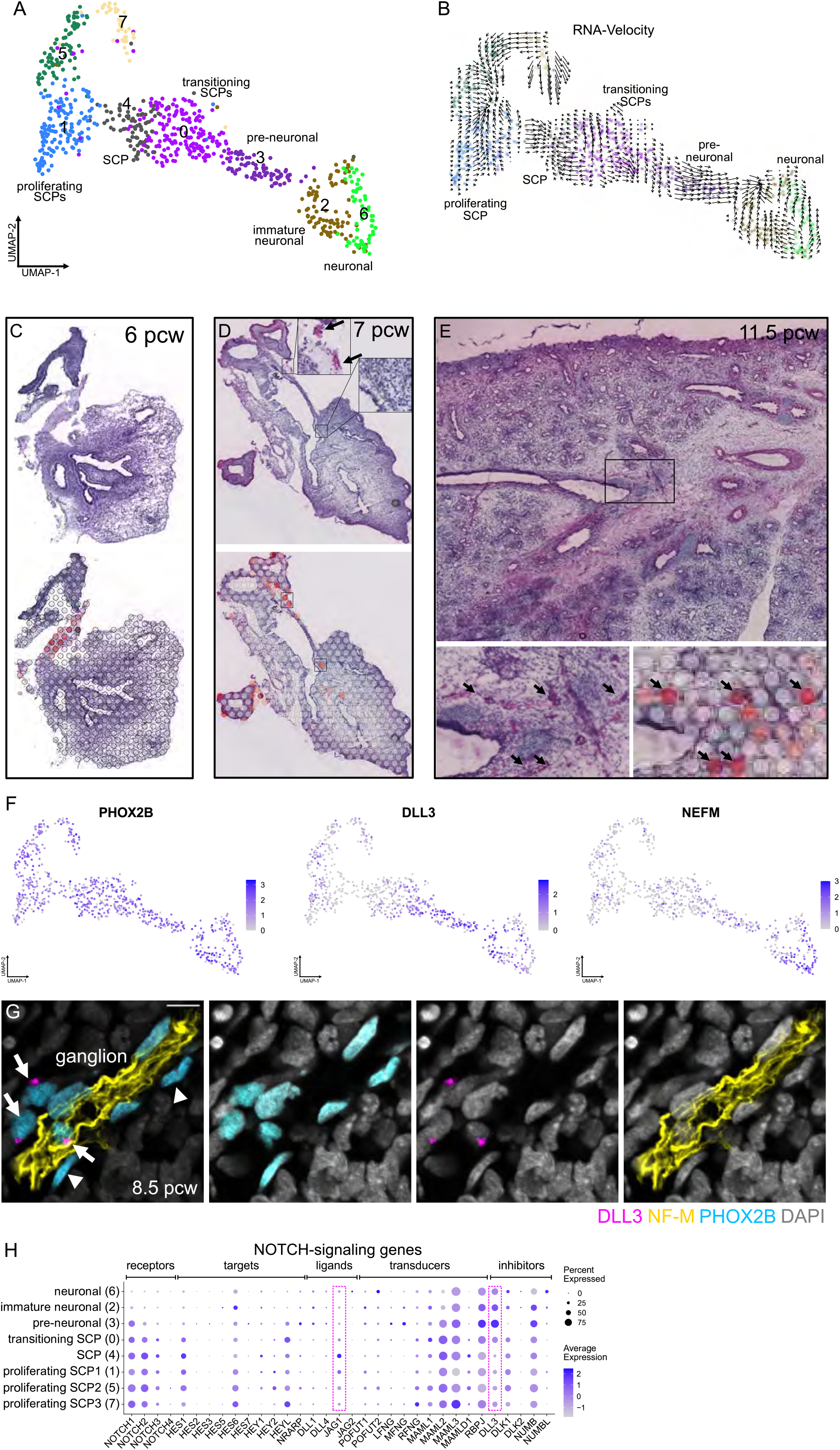
Parasympathetic neuron trajectory in the embryonic lung. **(A)** UMAP-plot of 752 neuronal cells. Colors: suggested clusters. **(B)** RNA velocity analysis on the UMAP-plot. Colors As in “A” and direction of arrows shows the future state of the cells. **(C-E)** Stereoscope neuronal score on 6 (C), 7 (D) and 11.5 (E) pcw lung sections. Top panels: high-resolution H&E images. Bottom panels: Stereoscope-score of neuronal cells, (SCPs and neurons, together). Arrows: possible ganglia with high percent of neuronal cells. Dark red: high, gray: zero percent. **(F)** UMAP-plots of *PHOX2B* (SCPs-neurons), *DLL3* (developing neurons) and *NEFM* (NF-M) (mature neurons). Expression levels correspond to log_2_(normalized UMI-counts+1) (library size, normalized to 10.000). **(G)** Immunofluorescence of the PHOX2B (cyan), DLL3 (magenta) and NF-M (yellow). Nuclei: DAPI (gray). Scale-bar: 20 µm. Arrows: PHOX2B^pos^ DLL3^pos^ neurons. Arrowheads: PHOX2B^pos^ DLL3^neg^ SCPs. **(H)** Balloon-plot of NOTCH-signaling genes expression in neuronal clusters, including receptors, targets, ligands, transducers and inhibitors (Ouadah et al., 2019). Brackets highlight JAG1 and DLL3.

Stereoscope analysis, detected the signature of the whole cluster (both SCPs and neuronal cells) in regions outside the lung lobes at 6 pcw (Fig. 3C). Intra-lobar signal was first detected close to the trachea at 7 pcw (Fig. 3D). At later timepoints the signal was detected more centrally, within the bronchovascular bundle interstitium (Dalpiaz *et al*., 2017), coinciding with distinct H&E staining (Fig. 3E). This suggests that SCPs, presumably deriving from the neural crest, enter the lung and mature to parasympathetic neurons in ganglia embedded in the bronchial interstitium. To explore the cellular composition and differentiation states in the proposed embryonic ganglia we first stained for PHOX2B, expected to be expressed in both SCPs and neurons, DLL3, which is selectively activated in differentiating neurons and NF-M to visualize neuronal projections (Fig. 3 F, G). We found several clusters of PHOX2B^pos^ cells with Neurofilament staining, suggesting that they contain maturing neurons at 8.5 pcw. Among the 8 PHOX2B^pos^ cells in a NF-M^pos^ domain, 3 expressed DLL3 suggesting that they correspond to differentiating neurons (mainly cl-2 and −3). We further explored this, by staining for the characteristic TFs SOX10, ASCL1 and ISL1, which are sequentially activated along the SCP-neuronal trajectory (Suppl. Fig. 3 C-E). We detected SOX10^pos^ SCPs, SOX10^pos^-ASCL1^pos^ neuronal precursors and ISL1^pos^ neurons, consistent with the differentiation steps proposed by pseudotime analysis (Suppl. Fig 3F). The selective expression of ASCL1 and DLL3 in subclusters of the ganglionic cells prompted us to interrogate the expression of NOTCH signaling pathway genes in the clusters (Fig. 3H). The selective expression of *JAG1* in SCPs suggested that it activates and maintains the SCP state and/or drives the formation of immature Schwann cells as proposed in studies of mouse limb nerves (Woodhoo et al., 2009).

The localization of parasympathetic ganglia in the bronchial interstitium and the NF-M positive projections in 8.5 pcw lungs suggested the early onset of signaling events leading to ASM innervation by parasympathetic post-ganglionic neurons (Burns *et al*., 2008; Freem *et al*., 2010). We analyzed possible interactions between the neurons and the ASM cell-identities focusing on ASM-derived signals with receptors and potential targets specifically expressed in neurons (Suppl. Fig. 3 G-J). The ASM-produced chemokine CXCL12 could activate its receptor CXCR4 in neurons (Suppl. Fig. 3I), regulating processes like the neuronal migration, as previously described for midbrain dopaminergic neurons (Yang et al., 2013). Similarly, THBS2 secreted by ASM is predicted to activate the neuronal migration through the CD47 receptor and induce the *TBX20* and *GFRA* targets, in mature neurons (Sergaki and Ibanez, 2017; Song et al., 2006) (Suppl. Fig. 3J). We conclude that pulmonary parasympathetic neurons derive from SCPs, similar to neurons in human sympathetic adrenal ganglia (Kameneva et al., 2021) and in mouse parasympathetic ganglia (Dyachuk *et al*., 2014; Espinosa-Medina *et al*., 2014). Lineage trajectories and spatial analysis suggest that NOTCH-mediated signaling, among SCPs and neurons, establishes and maintains ganglionic populations and that short- and long-range interactions of neurons with ASM may mediate bronchial innervation.

### Early developmental trajectories of epithelial differentiation

We subclustered epithelial cells into 15 groups (Fig. 4A) and annotated these based on characteristic markers (Suppl. Fig. 4A, B) and their spatial distribution (Fig. 4 B, C). We detected four distal epithelial identities (epi cl-10, −2, −3 and −9), with distinct positions and maturation states, seven proximal types, corresponding to ciliated (epi cl-14), secretory (epi cl-0), neuroendocrine (NE) cells (epi cl-11 and −12) and their progenitors (epi cl-6, −7 and −4). We also found an intermediately located *SOX2*^*medium*^ *SOX9*^*low*^ population (epi cl-1) and three proliferating cell-states characterized by *MKI67* and *PCNA* expression (epi cl-8, −13 and −5).

**Figure 4.**
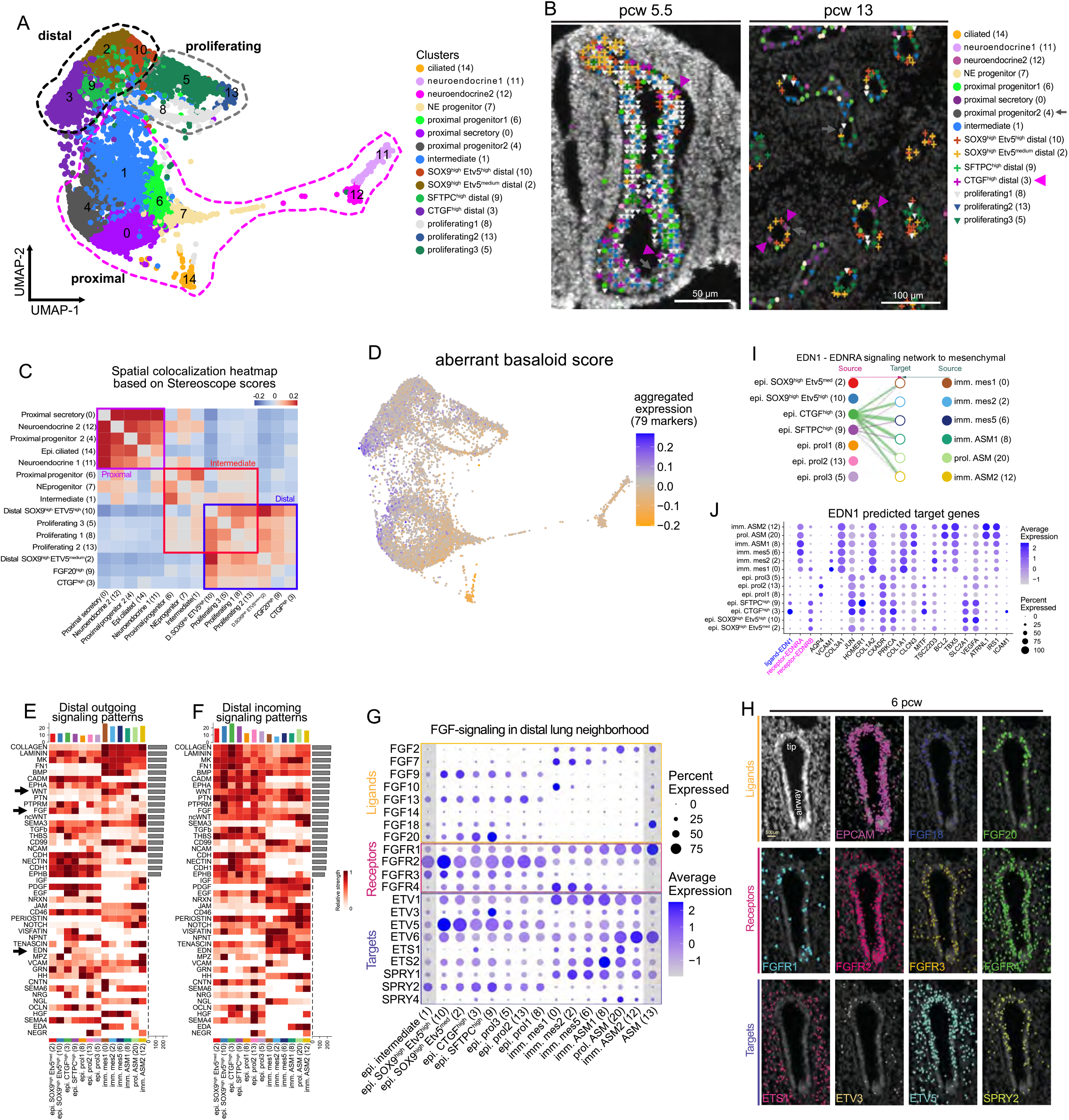
Epithelial diversity in developing human lungs. **(A)** UMAP-plot of 10940 epithelial cells. Colors: suggested clusters. Dotted outlines: main cell groups of proximal (magenta), proliferating (grey) and distal cells (black). **(B)** Annotation of segmented airway cell areas by PciSeq, using HybISS data in 5.5 pcw (left) and 13 pcw (right) airways. Colors as in “A”, distal clusters: cross, proliferating: inverted triangle and proximal: circle. Gray arrows: prox. progenitor2 (cl-4), magenta arrowheads: *CTGF*^high^ distal (cl-3). **(C)** Heatmap of the spatial correlation of the indicated clusters, based on Stereoscope scores (ST-data). Positive correlations: red, negative correlations: blue. Brackets: distal, intermediate and proximal main patterns. **(D)** UMAP-plot showing the activated-epithelial (basaloid) score, as indicated by the aggregate expression of basaloid cell (Adams *et al*., 2020) selective markers. **(E-F)** CellChat heatmaps of the significant outgoing (E) and incoming (F) signaling patterns in distal lung neighborhood (Fig. 1D). Bars represent the outgoing/incoming overall potential on each cluster (top) and pathway (right). **(G)** Balloon-plot of FGF-ligand, -receptor and -target expression levels, in distal lung clusters. Balloon size: percent of positive cells. Color intensity: scaled expression. Epithelial intermediate (cl-0) and mesenchymal ASM (cl-13): as control cell-states (not in the specific neighborhood, with gray shadow). **(H)** HybISS in situ validation of FGF-pathway genes, DAPI: nuclei (top-left). Top: general epithelial marker *EPCAM, FGF20* and *FGF18* ligands. Middle: *FGFR1-4* receptors. Bottom: *ETS1, ETV3, ETV5* and *SPRY2* targets. **(I)** CellChat-predicted, *EDN1*-*EDNRA* communication pattern in distal lung. **(J)** Balloon-plot of the top-20 *EDN1*-predicted target genes by NicheNet. Ligand: blue, Receptors: magenta.

Epithelial clusters −2, −3, −9 and −10, corresponding to distal epithelial cells, are characterized by *SOX9* (Chang et al., 2013; Nikolic *et al*., 2017) and *ETV5* expression (Suppl. Fig. 4A). Epithelial cl-2 and −10 show the highest *SOX9* levels and are located in the most distal part of the developing bud tips (Fig. 4 B, C). The PAGA-plot and their topology suggested that they may be the source of the remaining two distal SFTPC^pos^ clusters (Fig. 4B, Suppl. Fig. 4C, D), which are predominantly composed of cells from older than 10 pcw embryos (Suppl. Fig. 4E). Accordingly, distal epithelial cl-9 cells include biosynthetically more active cells, with higher *SFTPC* (Kalina et al., 1992) and *ACSL3* expression (Suppl. Fig. 4 F, G). *ACSL3* participates in lipid metabolism (Padanad et al., 2016), a perquisite for surfactant biosynthesis (Agassandian and Mallampalli, 2013). By contrast, epithelial cl-3 cells were found scattered in distal epithelium as early as 5 pcw (Fig. 4B) and express elevated levels of CTGF (Suppl. Fig. 4 A, H), a growth factor for mouse alveolar development (Baguma-Nibasheka and Kablar, 2008), which also stimulates lung fibroblasts, in fibrosis models (Yang et al., 2014). cl-3 cells also express KRT17 (Suppl. Fig. 4I), a characteristic marker for basaloid cells, a pathogenic cell-state in interstitial pulmonary fibrosis (IPF) (Adams *et al*., 2020). To further explore similarities of cl-3 and basaloid cells, we used the gene expression signature of selective basaloid genes (Adams *et al*., 2020) to score all epithelial cells of our dataset. We found that cells from cl-3, present in distal epithelium showed the highest expression score for 79 of the 85 selective markers (Fig. 4D), revealing a shared epithelial genetic program in lung development and fibrosis.

### Interactome analyses indicate dynamic communications within and between distal epithelial and mesenchymal cell-identities

We utilized the definitions of cell neighborhoods (Fig. 1D) to explore candidate pathways in cell-cell communications, in the distal lung compartment (Figure 4 E, F). FGF-signaling was among the most prominent predictions, but unlike the mouse embryonic lung, where mesenchymal Fgf10 induces branching through Fgfr2b, in the distal branch tips (Bellusci *et al*., 1997b), the predictions of FGF-sources, in the distal human lung, involved both mesenchymal and epithelial ligands. *FGF10* is expressed in mes cl-0 (Fig. 4G) and this cluster is found scattered in positions around the epithelium (Bellusci et al., 1997a; Danopoulos et al., 2019). Our HybISS analysis detected *FGF18* transcripts in 4 clusters (epi cl-2, −3, −9 and −10) and more selective *FGF20* expression in clusters cl-9 and cl-3 (Fig. 4H). The epithelial expression of *FGFR2, FGFR3, FGFR4* and the localized expression of potential downstream targets *ETV5* (Herriges et al., 2015) *SPRY2* (Mailleux et al., 2001), suggests an epithelial intrinsic/ autonomous function of FGF-signaling in the human lung (Fig. 4 G, H). A prominent predicted target of epithelial FGFR activation is *SOX9* (Suppl. Fig. 4J), consistent with experiments reporting its regulation by FGF/Kras signaling in embryonic lungs (Chang *et al*., 2013; Danopoulos *et al*., 2019).

Considering the prominence of the basaloid gene expression program in IPF pathophysiology (Adams *et al*., 2020; Strunz et al., 2020), we analyzed the communication patterns between cl-3 and the other cell-states of the distal lung neighborhood. Among the most contributing pathways, we found EDN-signaling, from cl-3 cells, targeting mainly the mesenchyme (Fig. 4 I, J). Interestingly, in the context of fibrotic disease, adenovirus-mediated *EDN1* overexpression by epithelial cells causes lung fibrosis (Lagares et al., 2012). Importantly, target-gene prediction analysis indicates that EDN1 can up-regulate the collagen genes, in addition to *TBX5* (Fig. 4F). This suggests that EDN1 signaling is involved in the communication of “activated” epithelial cells (cl-3) with the surrounding stromal cells, inducing not only ECM genes but also maintaining high levels of *TBX5* expression in mesenchymal progenitors, to facilitate normal branching morphogenesis (Arora et al., 2012).

### Distinct steps in proximal airway cell differentiation

Among the proximal epithelial clusters, secretory *SCGB3A2*^pos^ (Reynolds et al., 2002) cl-0 and cl-4, together with neuroendocrine *ASCL1*^pos^ (epi cl-11 and −12) and ciliated *FOXJ1*^pos^ (epi cl-14) were located in the most proximal locations, whereas the proximal progenitor (cl-6) and NE progenitor (cl-7) cells were found in slightly more distal positions (Fig. 4C). Epithelial cl-14 is characterized by *FOXJ1* and early ciliogenesis genes, suggesting that it contains immature ciliated cells (Suppl. Fig. 4K). Among the secretory clusters, the major difference between cl-0 and cl-4 was the high levels of *HOPX* and *KRT17* in cl-4 (Suppl. Fig. 4 A, B), which also showed a high score for the activated-epithelial/basaloid program (Fig. 4D), suggesting similarities between cl-4 and the distally localized epithelial cl-3. We did not detected differences in the spatial distribution of cl-0 and cl-4 cells (Fig. 4 B, C), but comparison of their gene expression profiles revealed that cl-4 is enriched for migration-related genes (Suppl. Fig. 4L). Thus, cl-4 may correspond to a transient airway secretory progenitor state giving rise to the “default”, static airway secretory cells of cl-0 (Suppl. Fig. 4C). PAGA-plot (Suppl. Fig. 4C) and Velocyto (Suppl. Fig. 5A) analyses suggested that the immature epithelial cl-6 can function as a source for the secretory cl-0 and the progenitor epithelial cl-7, which further progresses to the neuroendocrine epithelial cl-11 and cl-12 (Fig. 5A). Pseudotime analysis along the two trajectories identified 569 differentially expressed genes that were clustered in 9 statistically significant modules (Suppl. Fig. 5B). Among the earliest genes activated in the secretory trajectory we detected *YAP1* and *GPC5*, a WNT-signaling extracellular inhibitor (Fig. 5B, gene module-6) (Ostrin et al., 2018; Yuan et al., 2016). These were followed by increased levels of the characteristic secretory marker, *SCGB3A2* and the NOTCH-signaling targets, *HES1* and *HES4* (Fig. 5B, gene module-9), further arguing for its importance in airway secretory cell differentiation (Tsao et al., 2009) and maintenance (Lafkas et al., 2015).

**Figure 5.**
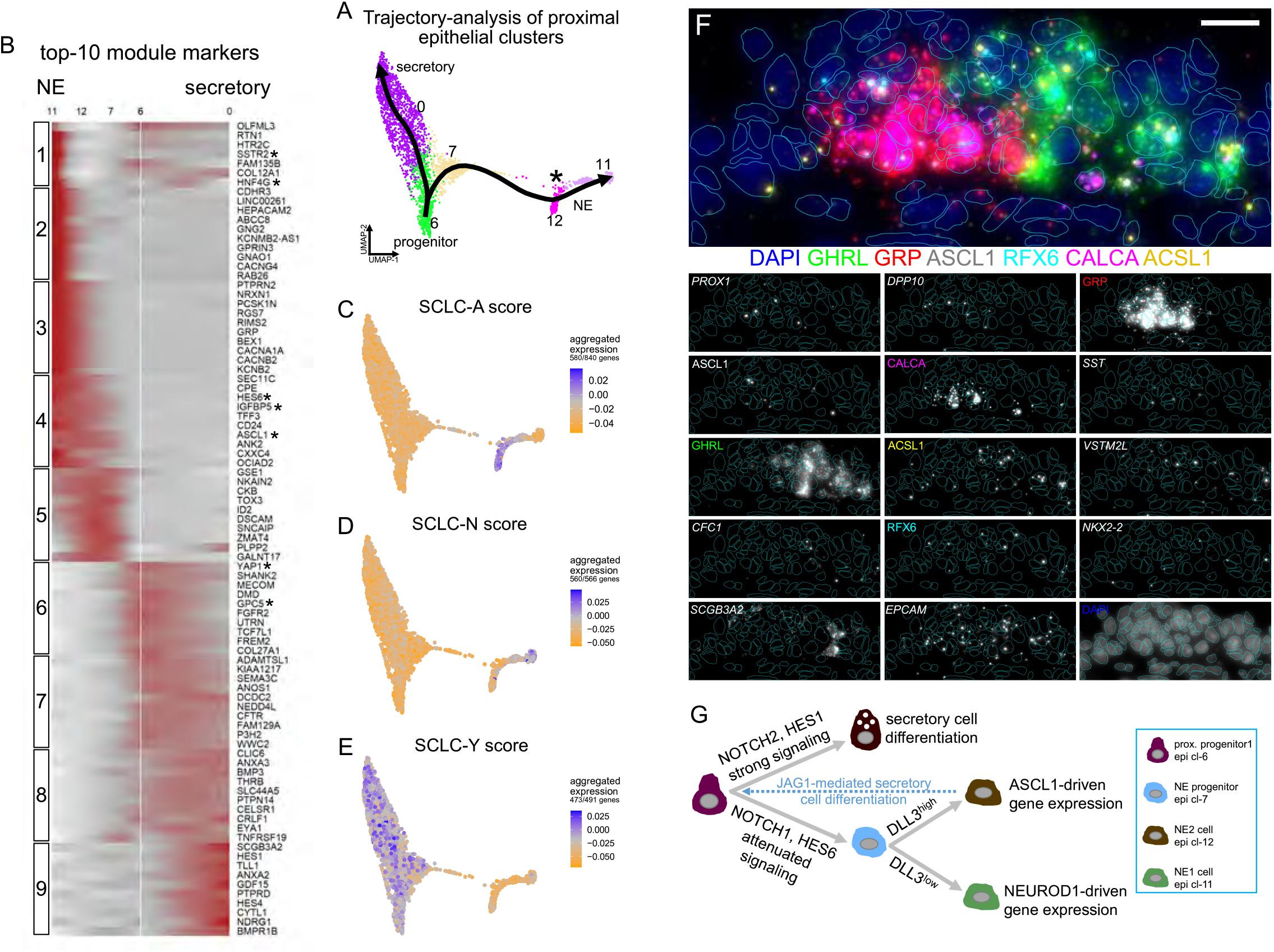
Analysis of proximal epithelium developmental trajectories. **(A)** UMAP-plot of proximal clusters and pseudotime of secretory and neuroendocrine (NE) trajectories, estimated by Slingshot. Colors as in Fig. 4A. The asterisk: bifurcation point of the 2 NE-clusters. **(B)** Heatmap of the top-10 markers of each stable gene-module of the 569 differentially expressed genes along the 2 trajectories (Suppl. Fig. 5B), according to tradeSeq. **(C-E)** UMAP-plots showing the aggregate expression of the (C) SCLC-A, (D) SCLC-N and (E) SCLC-Y signature-genes (Ireland *et al*., 2020). **(F)** SCRINSHOT-analysis of NE-clusters, in a 14 pcw proximal airway. Cl-11 markers: *GHRL* (green), *RFX6* (cyan), *ACSL1* (yellow). Cl-12 markers: *GRP* (red), *ASCL1* (gray), *CALCA* (magenta). Dotted outlines: manually segmented nuclei. *PROX1, DPP10, SST, VSTM2L, CFC, NKX2-2, SCGB3A2* and *EPCAM* (italics) only as individual channels. Scale-bar: 10µm. **(G)** Schematic representation of the suggested NOTCH-signaling function on secretory and NE-cell specification.

### Two neuroendocrine identities with distinct topologies and possible functions

In the neuroendocrine trajectory, epithelial cl-7 likely represents a NE-progenitor state expressing low levels of *ASCL1*, a critical factor in NE-cell development (Borges et al., 1997) (Fig. 5B: gene module-4). Analysis of all differentially expressed TFs along the secretory and NE trajectories (Suppl. Fig. 5C) showed that the direct ASCL1-target, *MYCL* (Borromeo et al., 2016) was transiently activated along the NE-trajectory (Suppl. Fig. 5 C, D), indicating a role in NE-cell specification. Epithelial cl-7 is connected with some cells with the NE cl-12, creating a stalk that splits into two directions, one towards the remaining cl-12 and the other towards cl-11 (Fig. 4A, asterisk in 5A). In this part of the trajectory, we identified, except for *ASCL1*, its direct target *IGFBP5* (Wang et al., 2019), together with *HES6*, (Nelson et al., 2009) (Fig. 5B, gene module-4). By contrast, the cl-11 NE-state was characterized by *NEUROD1* (Suppl. Fig. 5E), its target TF *HNF4G* (Borromeo *et al*., 2016) (Fig. 5B, gene module-1, Suppl. Fig. 5F) and the somatostatin receptor SSTR2 (Fig. 5B, gene module-1), resembling a recently identified, neuroendocrine cell-type in human embryos (Cao *et al*., 2020). We further compared cl-11 and cl-12 gene expression profiles (Suppl. Fig. 5G, Suppl. Table1). cl-12 produces the characteristic pulmonary neuropeptides GRP and CALCA together with SST, whereas cl-11 expressed *GHRL* and *CRH*. GO-analysis for enriched biological processes in cl-11 suggests hormone secretion (GO:0030072) and neuronal axon guidance (GO:0007411), as characteristic terms (Suppl. Fig. 5 H, I). The expression of *ASCL1* in cl-12 and *NEUROD1* in cl-11, suggested that these NE-identities resemble distinct states in Small Cell Lung Carcinoma (SCLC) progression. We used the gene-expression signatures of the SCLC-A (ASCL1^high^), SCLC-N (NEUROD1^high^) and SCLC-Y (YAP1^high^) cancer clusters (Borromeo *et al*., 2016; Ireland et al., 2020) to score similarity with the clusters in the NE-trajectory of our dataset. This showed that the SCLC-A signature is shared with cl-12 (Fig. 4C), the SCLC-N with cl-11 (Fig. 4D) and the SCLC-Y with the cl-7, −6 and −0 (Fig. 4E). This suggests that the embryonic pseudotime direction of NE-differentiation, from naïve secretory cells, is reversed during SCLC progression.

To investigate the spatial relation of the two NE-clusters, we selected a panel of 31 genes. These included: 1) general NE-markers *(PROX1, DPP10*), 2) cl-12 selective genes (*ASCL1, GRP, SST* and *CALCA*), 3) cl-11 markers (*GHRL, ACSL1, RFX6, ARX, CFC1, VSTM2L, PCSK1* and *NKX2-2*), and 4) markers of the major epithelial and mesenchymal clusters (Fig. 5F, Suppl. Fig. 5 J-L). To identify neuroendocrine-specific patterns, we segmented the sections in hexagonal bins with a 7 m width, approximating the size of individual cells. By exploring the heterogeneity of 20351 bins expressing general epithelial and characteristic NE genes (STAR Methods), we confirmed the presence of three main NE-associated cell types, corresponding to NE progenitors, GRP^pos^ and GHRL^pos^ NE-cells *in situ* (Suppl. Fig. 6 A, B). The expression patterns of these populations match the ones observed in our scRNA-seq dataset. Interestingly, GHRL^pos^ NE-cells were located exclusively in most proximal airways, while NE progenitors and GRP^pos^ NE-cells were less restricted in their location along the airway proximal-distal axis (Suppl. Fig. 6 C-F). As expected, the relative frequency of neuroendocrine progenitor cells was found to decrease through the different developmental stages, in contrast with mature neuroendocrine cells (Suppl. Fig 6G).

Because different levels of graded NOTCH-signaling activation are required for the transition of SCLC-A cells to SCLC-N and SCLC-Y (Ireland *et al*., 2020), we interrogated the clusters in the embryonic NE-pseudotime trajectory for NOTCH-signaling genes (Suppl. Fig. 5M). The cl-12 NE-cells express high levels of *JAG1* and of the cell-autonomous inhibitor *DLL3* (Ladi et al., 2005), in addition to low levels of *JAG2* and *DLL1* ligands. The pathway target and inhibitor *HES6* (Nelson *et al*., 2009) was expressed by both NE-clusters. This suggests that cl-12 cells are the source of NOTCH-signaling in their neighborhood but they are less capable of receiving it. The downregulation of *DLL3*, in some cl-12 cells, activates NOTCH-signaling and initiates the cl-11 gene-expression program defined by the *NEUROD1, RFX6, HNF4G* and *NKX2-2* TFs (Suppl. Fig. 5 C, E, F). More upstream in the trajectory, at the bifurcation of secretory (cl-6) and NE-progenitor (cl-7) states, the repressor *REST* (Lim et al., 2017) and the receptor *NOTCH2* showed similar expression levels but *HES6* and *NOTCH1* were stronger expressed in the NE-progenitor cluster, suggesting differences in the strength and duration of NOTCH-signaling (Liu et al., 2015; Liu et al., 2013). In particular, NOTCH2-mediated activation in proximal progenitors (cl-6) is expected to be more potent, promoting the secretory differentiation. Overall, the pseudotime analysis suggests two sequential but distinct NOTCH-signaling events, utilizing different ligands and intracellular effectors, one to promote secretory differentiation and the other to inhibit cl-12 cells from acquiring the cl-11 state (Fig. 5G). Further interactome analysis revealed another unique communication pattern between the two NE cell-identities involving somatostatin (SST) from cl-12 and its receptor SSTR2 in cl-11 (Suppl. Fig. 5N). We found the anti-apoptotic factors BCL2 (Singh et al., 2019), CITED2 (Mattes et al., 2017) and IRF2BP2 (Pastor et al., 2021) (Suppl. Fig. 5O) among the top-100 predicted target-genes arguing that the cl-12 cells prevent apoptosis of the cl-11 cells, through SST. This is consistent with the decelerating effect of SSTR2 inhibition on SCLC growth in vivo and in vitro models (Lehman et al., 2019).

In summary, we mapped the distinct topologies and developmental trajectories of two NE-identities (Cao *et al*., 2020) from naïve epithelial cells in the embryonic lung. Each trajectory contains distinct candidate regulators of NOTCH-signaling for the respective cell-state transitions. The embryonic NE-cells share gene expression programs with SCLC subtypes and might function as a reference to better understand the aberrant transitions of cancer cells.

### Zonation patterns of mesenchymal progenitors along two epithelial axes

Stromal cell populations in fully-grown lungs show distinct distributions along the proximal-distal axis of the airway tubes. They also show specialized radial arrangements surrounding each major airway, with ASM cells adjacent to the epithelium (center) and adventitial fibroblasts and chondroblasts positioned more peripherally. To explore the spatial organization of different mesenchymal trajectories (AF, ASM and adventitial fibroblast) relative to the growing airway branches on the tissue level, we defined two axes. A proximal-distal one, along the airway length starting from the root of the branch and a radial extending from the airway center towards peripheral positions in the mesenchyme. We positioned the ST-spots and the cells detected using ISS relative to these two airway-dependent axes (Fig. 6A, STAR Methods). Analysis of tissues from different stages by the two spatial methods for immature and more differentiated states of adventitial fibroblasts (mes cl-10), ASM (mes cl-13) and AFs (mes cl-16) (Fig. 6A, B, Suppl. Fig. 7) revealed that more immature cell states occupy predominantly distal peripheral positions. By contrast, the more mature ones are more proximal and generally located centrally. The most immature ASM clusters (cl-0, −2 and −6) were the most peripheral. More differentiated clusters (cl-8, −20 and −12) were found closer to the airways and in more proximal positions, whereas the most mature ASM (cl-13) was found proximal and tightly associated with the airways. At all three consecutive time points (6, 8.5 and 11.5 pcw), the immature fibroblast (mes cl-4) was consistently found more proximal compared the ASM progenitor clusters. This argues for the presence of a peripheral central zone of mesenchymal progenitors giving rise to adventitial fibroblasts (AdvF), airway fibroblasts (AFs) and chondroblasts (CBs) and reveals an early origin of radial patterning in the mesoderm. We suggest that undifferentiated cells from the distinct progenitor regions are expected to proliferate and continuously differentiate while migrating radially towards the center and their functional positions.

**Figure 6.**
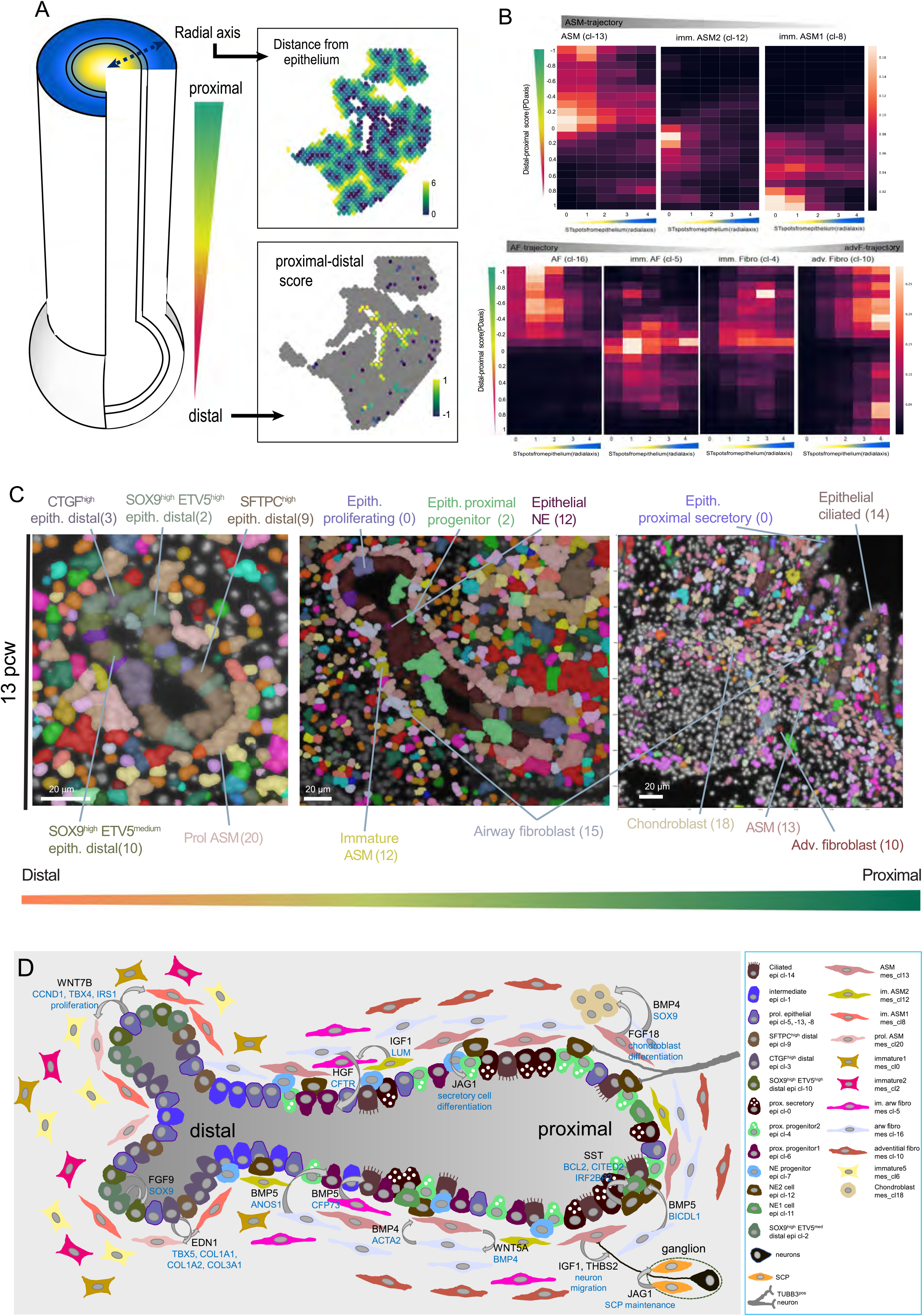
Assessing the molecular complexity of embryonic human airways. **(A)** (left) Schematic representation of the radial (distance from epithelium) and proximal-distal airway-dependent axes. (right) Spatial maps of the radial (top) and proximal-distal (bottom), scores of an 8.5 pcw section, analyzed by ST. **(B)** Heatmaps of ASM, AF and Adventitial fibroblast-related cluster cell-densities in the two analyzed axes. **(C)** Spatial cell-type maps of distal (left), intermediate (middle) and proximal (right) airways. Segmented nuclei are colored according to the most probable, predicted cell-type. Colors as in Fig. 1B. **(D)** Scheme of the cellular and molecular complexity in developing lung. The included cell-types were identified via scRNA-Seq and their spatial context was defined by spatial methods. CellChat-predicted communication patterns: curved arrows. NicheNet-predicted ligands (black) and corresponding target genes or outcome: cyan text. (right) Description of all involved cell-types and sensory neurons (not found in the scRNA-Seq).

### Synopsis of cellular heterogeneity and possible cell communication patterns during the first trimester

The utilized probabilistic methods [PciSeq (Qian et al., 2020) and Tangram (Biancalani et al., 2020)] facilitated the creation of systematic spatial maps of several developmental stages, showing the cellular composition of distinct organ compartments over time (Fig. 6C). On the tissue level, this allows to define spatial rules of tissue organization and to estimate developmental origins by interrogating the relative positions of pseudotime trajectories. A graphical representation of the developing lung, shows a summary of mature and intermediate cell states, localized in distinct tissue positions, creating cell “neighborhoods”, which are defined by specific communication patterns (Fig. 6D). We have produced an online platform for direct access to the ISS, ST and SCRINSHOT spatial analyses, together with the scRNA-Seq analysis and CellChat results via the TissUUmaps viewing tool (Solorzano et al., 2020) (https://tissuumaps.dckube.scilifelab.se/web/private/HDCA/index.html). This site provides an easily accessible and interactive atlas of early lung development without the need of downloading image data or installing software to directly facilitate exploration, sharing and hypothesis building.

## Discussion

We have generated the first topographic atlas of the developing human lung, combining gene expression profiling by scRNA-Seq with spatial transcriptomics on intact tissues. We identified and mapped 83 cell-states and inferred developmental trajectories leading to a remarkable heterogeneity reflecting the structural and functional complexity of the lung. Although we present an extensive analysis of weekly intervals during the first trimester (a co-submitted paper focuses on the second trimester), our data have a few limitations. Our first datapoint is at 5 pcw and we only analyzed about 180 thousand cells. Earlier and broader sampling is likely to uncover additional diversity and infer more precise trajectories than the current ones. We aimed to collect and analyze freshly dissociated cells from whole lungs without enrichment for specific populations. The lack of enrichment may have hampered detection of rare, fragile or difficult to dissociate cells. Indeed, we detected chondroblasts and mesothelial cells only in the samples deriving from earlier timepoints. We performed iterative clustering, where a conservative first clustering was followed by sub-clustering of the major populations. Although most of the subclusters showed distinct topologies, some of the cell-states may result from over-clustering. Finally, we have described the spatial diversity of the developing lung only at the mRNA level, relating this diversity to the proteome and further to physiological functions remains a future task.

We suggest that the diversity of gene expression patterns in the developing human lung can be explained at distinct but hierarchically coupled levels. First, the major cell-classes of epithelial, endothelial, immune, stromal and neuronal cells are characterized by the distinct gene expression programs of their ancestries from distinct germ layers; endoderm, mesoderm and ectoderm. Several levels of subdivisions in each of these classes occur during the first trimester. For example, within the endothelial group there are lymphatic, venous, arterial, bronchial and capillary clusters characterized by distinct regulatory and functional gene-expression profiles. Second, some cell-clusters show region-specific gene expression profiles, presumably reflecting their developmental history. This is exemplified, by the separation of proximal and distal compartments in the epithelium. The SOX2-proximal and the SOX9-distal domains are specified earlier and are maintained during the glandular stages. This suggests that transcriptional networks are conveyed into the later diversification of more specialized cell states specific to each region. Our spatial analysis illustrates this by the striking correlation of characteristically different radial arrangements of airway fibroblasts and smooth muscle states along different positions of the epithelial proximal-distal axis. This suggests that the different values of the proximal-distal axis intersect with distinct values of a radial axis visualized by the organization of surrounding smooth muscle and fibroblast states. The potential regulatory relationships between these axes are unknown. A third level of diversification results from cell interactions within the local environments reflecting inducible or transient activation or repression of gene-modules to signals. The integration of single-cell sequencing with spatial transcriptomics data defined specific neighborhoods for most of the cell-states. Our curated interactome analysis predicted several known and new examples of this organization level. They include the activation of NOTCH-signaling between the SCP and neuronal states, promoting neuronal differentiation in parasympathetic ganglia and the potential chemokine signaling pathway that potentially controls ASM innervation.

Lung diseases are major causes of death worldwide (Gibson et al., 2013). An outstanding challenge for medical research is to define deviation points from normal cellular trajectories at the start and during the advancement of lung pathologies and to analyze cellular responses after treatments (Rajewsky et al., 2021). Our spatial analysis in the developing lung revealed several distinct cell-states, their interactions with neighbors and progression along differentiation trajectories. Comparison of these with the few published lung disease trajectories demonstrates the medical relevance of our approach. We found a striking reversal of the gene expression profiles of our trajectory from naïve epithelial cells to secretory and neuroendocrine states, in the progressively aggressive states of SCLC. The neighborhood-based interactome analysis showed that SST-signaling is only employed locally in the communication between the NEUROD1^pos^ (immature) and ASCL1^pos^ (mature) NE-clusters, suggesting that its inhibition may only affect the SCLC-A to SCLC-N transition. Similarly, the proposed differentiation path of migratory secretory cells in the distal epithelium, recapitulates the genetic program of basaloid cells, activated in fibrotic lungs and identifies the HBEGF and EDN1 signaling pathways as potential local regulators of mesenchymal proliferation and ECM-protein secretion. As single cell analysis technologies are increasingly used in the description of derailed cell-state trajectories in disease, we believe that our integrated scRNA-Seq data, with spatial transcriptomics and local interactome analysis in an open, interactive viewer will provide a useful resource towards understanding and reversal of pulmonary disease progression.

## Experimental Procedures

### Human lungs

Use of human fetal material from elective routine abortions was approved by the Swedish National Board of Health and Welfare and the analysis using this material was approved by the Swedish Ethical Review Authority (2018/769-31). After the clinical staff acquired informed written consent by the patient, the retrieved tissue was transferred to the research prenatal material. The lung sample was retrieved from fetuses between 5- and 14-weeks post conception (pcw).

### Tissue treatment for spatial analyses

One of the two lungs (preferentially the left), from each donor, was snap frozen in cryomatrix and further used for histological analyses. We cut 10-12μm-thick tissue sections with a cryostat (Leica CM3050S or analogue) and collected them onto poly-lysine coated slides (VWR Cat No. 631-0107) for SCRINSHOT and immunofluorescence or superfrost-plus (VWR Cat No. 48311-703) for ISS. Sections were left to dry in a container with silica gel or at 37°C, for 15 minutes and then stored at −80°C until usage.

### Tissue dissociation of human embryonic lungs

For tissue dissociation, lungs were finely minced with blades and micro scissors. For older timepoints, they were first dissected into smaller pieces. Then, they were digested in 4U/ml Elastase (Worthington, cat no. LS002292), 1 mg/ml of DNase (Worthington, cat no. LK003170) in HBSS (Gibco, cat no. 14170) at 37 °C ranging between 30 min to 3 h depending on the age (older timepoints require longer digestion times). Note, HBSS with 2 % FCS (Gibco, cat no. 10500064) was used for the whole procedure. The tissues were triturated with fire polished glass Pasteur pipettes every 15-20 min for the tissue to fall apart more easily and enhance the dissociation. After digestion, the cell suspension was filtered (twice if needed) in a 15ml Falcon tube using a 30μm cell strainer (CellTrics, Sysmex), to remove remaining undissociated clumps and debris. The filtered cell suspension was kept ice cold and was diluted (roughly 1:2) with ice cold HBSS to prevent further digestion by the enzyme. The filtered cell suspension was pelleted through centrifugation at 200g for 5 min at 4 °C, the supernatant was removed and the pellet resuspended in a small volume of calcium- and magnesium-free HBSS (Gibco, cat no. 14170) (starting with 100μl depending on age and size of the pellet). For erythrocyte and debris removal, if present, cells were layered on top of 3ml HBSS with 2 % FCS, in a 15ml Falcon tube and subjected to an additional centrifugation step at 100 g for 5 min at 4 °C. Next, cells resuspended in HBSS and transferred to 1.5ml Eppendorf tubes precoated with 30% BSA (A9576, Sigma-Aldrich). A Bürker chamber was used for cell counting.

### scRNA-seq of human embryonic lung cells

Single-cell RNAseq was carried out with the droplet-based platform Chromium Single Cell 3’ Reagent Kit v2 and v3. Cell suspensions were counted and diluted to concentrations of 800 – 1200 cells/μl for a target recovery of 5000 cells on the Chromium platform. Downstream procedures including cDNA synthesis, library preparation and sequencing were performed according to the manufacturer’s instruction (10X Genomics Inc.). Libraries were sequenced on an Illumina NovaSeq 6000 (Illumina). For the v2 and v3 libraries 75.000 and 200.000 reads/cell, respectively were targeted for sequencing. Across all 39 libraries, an average of 187242 reads/cell was obtained. Reads were aligned to the human reference genome GRCh38-3.0.0 and libraries were demultiplexed and aligned with the 10X Genomics pipeline CellRanger (version 3.0.2). Loom files were generated for each sample by running Velocyto (0.17.17) in order to map molecules to unspliced and spliced transcripts.

### Bioinformatic analysis for scRNA-Seq

All *.loom files were imported to R as “Seurat objects”, using the “connect” function of the loomR package and the “as.Seurat” function of SeuratDisk for *.loom files >3.0.0 (Hao et al., 2021; Stuart et al., 2019). The “spliced”, “unspliced” and “ambiguous” counts were obtained using the “ReadVelocity” function of SeuratWrappers package and we created objects with “merged”, “spliced”, “unspliced” and “ambiguous” counts.

The single-cell RNA sequencing (scRNA-Seq) datasets from the same donor that were sequenced in the same sequencing run were merged in order to create donor-specific objects. The only exception was the cells of donor17 that were analyzed as two individual datasets because 10×256 was sequenced after 10×253 but we identified no “batch effect” separating its cells from the others of the same donor (“10×253” and “10×256” in Viewer).

The individual donor datasets were analyzed separately using Seurat package in R, to inspect their quality. Firstly, we removed the cells with low and high number of detected genes, based on their histogram distribution, because they would correspond to cell fragments and multiplets, respectively. Next, we ran the DoubletFinder package (McGinnis et al., 2019) to identify and remove possibly cell-multiplets, considering that 4% of the analyzed cells are multiplets.

To integrate the resulting datasets of 163K cells, we used the SCTranform function in Seurat, with 5000 variable genes. We used 5000 integration features for the dataset integration, setting as reference dataset the donor17 that corresponds to the oldest time point of our analysis (14^th^ week post conception). Different integration approaches, except for using a reference dataset, produced errors because of the size of the whole dataset. We observed no profound clustering of the cells according the examined technical covariates, like the utilized 10X Genomics chemistry and the donor identity, especially for those of the same age (Viewer).

The principal component analysis (PCA) was based on the first 100 top principal components (PCs). For definition of the neighborhood graph and the clusters, we used the default settings of “FindNeighbors” and “FindClusters” functions of Seurat (Hao *et al*., 2021; Stuart *et al*., 2019), with 100 dimensions. For identification of selective markers for the proposed clusters we used the “FindAllMarkers” function (Hao *et al*., 2021; Stuart *et al*., 2019), with MAST (Finak et al., 2015) statistical test and maximum number of cells per cluster being set to 126, that corresponds to the smallest suggested cluster of the dataset. To accept a gene as a cluster marker, it had to be expressed in at least 25% of the cells in the cluster, have 0.1 logarithmic fold increase and be expressed in at least 10% more cell in the cluster than the remaining dataset. We also selected the statistically significant markers (adjusted p-value <0.001) for all downstream analyses.

For the analysis of (i) epithelial, (ii) endothelial and (iii) immune cells, we selected the corresponding clusters of the 163K cell dataset and harmonized the cells according to the donor-Parameter, Using the “PrepSCTIntegration” function in Seurat with default settings and 5000 features (genes) and regressing out stress-related genes (“AddModuleScore” function in Seurat) (Hao *et al*., 2021; Stuart *et al*., 2019), that have been previously shown to get induced by enzymatic tissue dissociation at 37°C (Denisenko et al., 2020). Because of the large size of mesenchymal cell-subset (>138K cells), we used donor17, as a reference dataset for the harmonization of the different donor datasets. Especially for the analysis of the neuronal cells, we selected the donor datasets with more than 29 cells, that facilitated their decent integration (5 pcw: 49 cells, 5.5 pcw: 187 cells, 6 pcw: 169 cells, 7 pcw: 227 cells, 8 pcw: 38 cells, 8.5 pcw: 52 cells and 14 pcw: 30 cells). The selected 752 cells were further processed as the other cell categories.

For dimension reduction and clustering of the above main cell-type categories, we applied the same approach as with whole dataset. But we used the first 50 PCs, because these cell-groups are relative homogeneous and a very large number of PCs added noise.

To further filter the cells for possible multiplets, we firstly normalized the counts to 10.000 and then we removed possible red-blood contaminants, setting expression of HBA1 < 4, when necessary. For each of the epithelial, endothelial and immune datasets, we detected a cluster that expressed mesenchymal cell markers. Taking into account that (a) mesenchymal cell number is 12-times larger than epithelial, 21-times than endothelial and 33-times than immune and (b) it is unlikely for immune cells to express mesenchymal cells markers, we considered these clusters, doublets and removed them.

For trajectory inference analysis of complex multi-cellular developmental tissue architecture, we guided our analysis towards understanding key lineage branching points inspired by the graph abstraction concept (Wolf *et al*.). We used the cell-cell unweighted shared nearest neighbour graph (SNN) (*G* ∈ {0,1} *N* × *N*) and their assigned one-hot clusters (*O* ∈{0,1} *N* × *k*) to compute for each cluster *k* the number of edges shared with all clusters (*E* ∈ ℜ *k* × *k*), including itself.

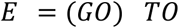

The number of cluster shared edges was then element-wise normalized by its total number of edges (Hadamard division), resulting in transition probabilities (*P*∈[0,1] *k* × *k*), that ranges 0 and 1 for each cluster, representing the proportion of connections shared between each cluster, where *J* ∈{1} *k* × *k*is a square all-ones matrix.

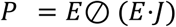

Spurious weak connections with transition probabilities below 10^−4^ were filtered out by setting its value to zero. Edges were then projected onto the cluster centroids on the UMAP embedding for visualization. Cluster transition probabilities on existing edges (*p ij*>0) were converted to graph weights (*w ij*) defined by the inverse of transition probabilities:

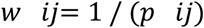

and optimal paths from immature (a.k.a. root). (a.k.a. root) to mature cells states were calculated (Csardi using dijkstra’s shortest path algorithm implemented in the igraph package(Csardi and Nepusz, 2006). The indicated clusters, for distinct trajectories, were selected and reanalysed to create a new UMAP-plot with “RunUMAP” function in Seurat (Hao *et al*., 2021; Stuart et *al*., 2019). The Slingshot package (Street et *al*., 2018) was used to do pseudotime analysis, along the trajectories. Firstly, we set the root and the end-point clusters with “getLineages” function and then we calculated the principal curves (“getCurves” function), the pseudotime estimates (“slingPseudotime” function) and the lineage assignment weights (“slingCurveWeights” function). To identify differentially expressed genes along the trajectories, we used the “fitGAM” function of tradeSeq package [6]. We used the “patternTest” function for the analyses of 2 trajectories with a common root and the “associationTest” function for the differential expression analysis along one trajectory. The cells were ordered according to pseudotime estimates and the differentially expressed genes were ordered based on the hierarchical clustering ward.D2 method, using “hclust” function in fastcluster package (Müllner, 2011) and plotted using a custom script. To identify modules of genes with the same expression pattern and their optimal number, we based on hierarchical clustering to define distinct number of clusters and used the “clusterboot” function of fcp package (Hennig and Imports, 2015) to calculate their stability values. For Velocyto analysis (La Manno et al., 2018), we used the “RunVelocity” function of SeuratWrappers.

For the analyses of aberrant basaloid (Adams *et al*., 2020) and SCLC (Ireland *et al*., 2020) gene expression programs in the scRNA-Seq dataset, we used the “AddModuleScore” function in Seurat (Hao *et al*., 2021; Stuart *et al*., 2019) to calculate the aggregated gene-expression scores of their characteristic markers, as they have been defined in the corresponding studies.

For the identification of transcription factors and co-factors, between the differentially expressed genes, we used the AnimalTFDB 3.0 database (Hu et al., 2019). The Human Protein Atlas was used for screening of secreted and surface (CD) proteins (Sjostedt et al., 2020; Uhlen et al., 2019) and Neuropedia database was used to find differentially expressed neuropeptides (Kim et al., 2011). Statistically significant (adjusted p-value <0.001, average logarithmic fold change >0.25) genes were used in Toppgene suite (Chen et al., 2009), for gene ontology (GO) analyses, with default settings.

## Spatial Transcriptomics (ST)

### ST library preparation

Spatial Gene Expression libraries (n=9) (6-13 pcw) were generated with the Visium Spatial Gene Expression Slide & Reagent kit (PN-1000184; 10X Genomics Inc.), according to manufacturer’s protocol. Prior to the analyses, RIN values were obtained for all samples to assess the quality of the RNA.

Depending on the size of each section, one or more sections of the same sample were placed in each capture area (6.5 × 6.5 mm) of the Visium arrays. The sections were first fixed for 10 min in acetone, stained with Mayer’ s Hematoxylin and Eosin Y and imaged with a Zeiss Imager.Z2 Microscope (Carl Zeiss Microscopy GmbH), using the Metafer5 software MetaSystems Hard & Software GmbH). Depending on the age of the lung, the tissue sections were permeabilized for 8 to 20 min to capture the mRNA molecules. The optimal fixative and permeabilization time for developing lung samples was determined prior to the Visium experiments using a Visium Spatial Tissue Optimization Slide & Reagent Kit (PN1000193; 10x Genomics Inc.). The cDNA synthesis and library preparation were done according to manufacturer’s protocol (PN-1000184, PN-1000215; 10X Genomics Inc.). Sufficient amount of 2nM to 4nM concentration libraries was used for sequencing for Illumina, Inc. platform, following manufacturer’s instructions.

### ST data analysis

Sequenced ST libraries were processed using Space Ranger 1.0.0 Pipeline (10X Genomics). Reads were aligned to the human reference genome to obtain an expression matrix. The count matrix was filtered for all mitochondrial, ribosomal and non-coding genes. Spots with less than 300 UMI, less than 100 genes and genes detected in less than 5 spots were excluded from the analysis. After filtering, a total of 18125 features were retained for final analysis across 66626 spots (6 pcw: 1439, 7 pcw: 2692, 8 pcw: 1840, 8.5 pcw: 1882, 9 pcw: 3284, 10 pcw: 11720, 11pcw: 15534, 12 pcw: 13287 and 13 pcw: 14948).

Normalization and dimension reduction were performed jointly using the Seurat and STUtility packages (version 0.1.0, https://ludvigla.github.io/STUtility_web_site/Installation.html). Technical variability across samples was reduced with RunSCT and RunHarmony (version 1.0, https://github.com/immunogenomics/harmony) functions. Principal Component Analysis (PCA) was used to select significant components and a total of 30 principal components were used in downstream analyses, in all cases.

### Integration of scRNA-Seq and ST data

For the integration between scRNA-Seq and Visium data, we used the Python package stereoscope (v.03) (Andersson *et al*., 2020). Raw counts from the scRNA-Seq and Visium data were used as input, along with the scRNA-Seq cluster labels. For the scRNA-Seq data from each donor, we used the top 5000 most variable genes as input, obtained by the “VariableFeatures” function in Seurat (Hao *et al*., 2021; Stuart *et al*., 2019). Stereoscope was run with 25,000 epochs with default parameters (more details in the “README” file in package github page). For the integrated scRNA-Seq, i.e. all age groups, the entire set of scRNA-Seq was used as input to each Visium sample individually and Stereoscope was run with 20,000 epochs. For visualization, the output matrix was imported into R and the stereoscope proportion values for each capture spot were plotted as features with the *STUtility* R package (v.1.0) (Bergenstrahle et al., 2020).

### Interactome analyses of spatially related cell-identities

For the definition of cell neighborhoods, that include cell-identities being consistently found with high percent in the same ST-spots, we used the stereoscope data and performed Pearson correlation analysis comparing the frequencies of the different cell-types in the analyzed ST-spots, across all samples and time points. We further proceeded with the pairwise connections, that had Pearson’s *r* higher than 0.04. The interactome analyses are based on CellChat (Jin *et al*., 2021) for the identification of ligand-receptor pairs that facilitate neighboring cell communications. We initially kept the genes with average gene expression >0.3 log_2_(normalized UMI-counts+1) in any of the analyzed clusters and then used default settings for the downstream analyses. To analyze the predicted target genes of specific ligands, we used the ligand-target score matrix of NicheNet (Browaeys *et al*., 2020) and selected the same genes as for CellChat, applying an extra filter by keeping the expressed in at least 25% of any of the clusters and have 10% increase in the number of positive cells and in the logarithmic fold-change. Then, we used Seurat to plot the 20 or 100 top-predicted genes, using “Dotplot” function. The ligand and the identified by CellChat receptors were also included at the beginning of the plot.

### In Situ Sequencing (HybISS)

#### Gene panel selection

The HybISS gene panel was selected based on two independent criteria: gene potential to be markers of the different identified populations and their role in different key signaling-pathways. To select the minimum amount of marker genes needed to uncover the cell-type of every cell in the analyzed samples, an initial list of candidate marker genes was generated by selecting the top 4 markers of the main clusters found when analyzing individually four samples from different timepoints (5 pcw, 8.5 pcw, 13 pcw and 14 pcw), based on their δct (difference in the percent of positives in the cluster against all other cells). This list was curated by assessing the importance of every gene in accurately predicting the different cell-types (https://github.com/Moldia/Tools/tree/master/Gene_selection). For this, ISS datasets were simulated by randomly distributing cells in a bidimensional space, assigning a cell-type to each cell and simulating the expression of each gene by sampling in a negative binomial distribution with r being the mean expression of a certain gene in a certain cell-type. Then, probabilistic cell-typing by ISS (pciSeq) (Qian *et al*., 2020) was used to assess the cell-type of each simulated cell, obtaining the contribution of each gene to predict correctly each cell-type. Top-5 genes contributing to correctly predict each cell-type were kept and further simulations were run, obtaining a final list of 72 genes that were able to predict correctly all the cell-types on simulated datasets. For the pathway gene selection, we interrogated the above four scRNA-Seq datasets for the expression of WNT, SHH, NOTCH and RTK pathway components, such as ligands, receptors, transducers, inhibitors and targets. We further processed with those that shown non-ubiquitous expression patterns. The final gene panel of 147 markers was sent to CARTANA with accompanying customized ID sequences for in-house HybISS chemistry detection.

#### HybISS mRNA detection

The HybISS experiments were performed by the ISS facility at Science for Life Laboratories (SciLifeLab) following manufacturer’s instructions of CARTANA’s High-Sensitivity library preparation kit, using customized backbones, as described in (Lee et al., 2020). After fixation, the tissue sections were overnight incubated with the probe mix, in a hybridization buffer, followed by stringent washing. Then, they were incubated with ligation mix. After washes, RCA was performed overnight. Finally, labelling for detection was performed as described in <protocols.io> (dx.doi.org/10.17504/ protocols.io.xy4fpyw). Twelve detection cycles were performed on each sample to avoid optical crowding. Therefore, detected genes were divided in three groups and their four cycle-based barcode was detected in either detection cycles 1-4, 5-8 or 9-12.

#### Imaging of HybISS detection cycles

Imaging was performed using a Zeiss Axio Imager.Z2 epifluorescence microscope (Carl Zeiss Microscopy, GmbH), with a Zeiss Plan-Apochromat 20x/0.8 objective (Carl Zeiss Microscopy, GmbH, 420650-9901) and an automatic multi-slide stage (PILine, M-686K011) to allow re-call of coordinates for the regions of interest, facilitating repetitive cycle imaging. The system was equipped with a Lumencor® SPECTRA X light engine LED source (Lumencor, Inc.), having the 395/25, 438/29, 470/24, 555/28, 635/22 and 730/40 filter paddles. The filters, for wavelength separation, included the quad band Chroma 89402 (DAPI, Cy3, Cy5), the quad band Chroma 89403 (AlexaFluor750), and the single band Zeiss 38HE (AlexaFluor488). Images were obtained with an ORCA-Flash4.0 LT Plus sCMOS camera (2048 × 2048, 16-bit, Hamamatsu Photonics K. K.).

#### HybISS image processing

Imaging data was processed with an in-house pipeline based on MATLAB (https://github.com/Moldia/iss_starfish). Maximum intensity projection was performed on each field of view in order to obtain a 2-dimensional representation of each tile. Then, stitching of tiles was performed using a MATLAB implementation of MIST algorithm, obtaining, after exporting, different *.tiff images corresponding to each channel and round. Then, data was retiled and formatted to fit the Starfish required input. Since genes can be either detected in 1-4, 5-8 or 9-12 detection cycles, each group was then decoded independently. Using Starfish tools, individual tiles were registered across cycles and a top hat filter was applied on each channel in order to get rid of the background noise. Channel intensities were also normalized and spots were detected. Finally, decoding was performed on each tile using MetricDistance, obtaining the identity of all the detected RCA products.

#### HybISS data analysis

Two different yet complementary strategies were followed to characterize the cellular heterogeneity within the in situ sequencing datasets. Probabilistic cell-typing for in situ sequencing (PciSeq) (Qian *et al*., 2020) was performed in order to identify the identity of every cell in the tissue. For this, cells were segmented based on DAPI using a watershed segmentation and reads were assigned to cells as described in (Qian *et al*., 2020). In addition, Tangram (Biancalani *et al*., 2020) was used to couple the scRNA-Seq with the HybISS datasets. Gene expression imputation was performed as described in (Biancalani *et al*., 2020) In 5 pcw sections, where nuclear segmentation was not possible, hexagonal binning was used to segment the tissue. In this case, the expression of each hexagonal bin was used as input for probabilistic cell-typing and Tangram.

## SCRINSHOT

### Gene selection, Padlock probe design and mRNA detection

For spatial analysis of the two identified NE-cell identities, we used the highly expressed *GRP* and *GHRL*, for easy identification of epi cl-12 and epi cl-11, respectively. Then, we selected markers that are expressed in intermediate and low levels, focusing mainly on TFs, like the *ASCL1, RFX6, NKX2-2, ARX* and *PROX1*. Markers like the *SCGB3A2, FOXJ1* and *TP63* were used to identify the non-neuroendocrine cells. The *SCGB1A1, SFTPC, ETV5, FOXJ1, AGER, SOX2* and *SOX9* padlock probes were designed as in (Sountoulidis *et al*., 2020). For the rest, a unique barcode was inserted in the backbone of all probes that recognize the same mRNA, that allowed their detection by only one detection oligo, reducing significantly the cost (all sequences are found in Suppl. Table2). All the reactions were done according to the original SCRINSHOT protocol, except for an increase of the detection-oligo hybridization temperature to 30 °C.

### Imaging of SCRINSHOT signals on tissue sections

For signal acquisition we did 13 detection cycles, using a Zeiss Axio Observer Z.2 fluorescent microscope (Carl Zeiss Microscopy, GmbH) with a Colibri 7 led light source (Carl Zeiss Microscopy, GmbH, 423052-9770-000), equipped with a Zeiss 20X/0.75 Plan-Apochromat, a Zeiss AxioCam 506 Mono digital camera and an automated stage, that allowed imaging of the same regions in every cycle. For signal detection, we used the following Chroma filters: DAPI (49000), FITC (49003), Cy3 (49304), Cy5 (49307), Texas Red (49310) and Atto740 (49007).

### SCRINSHOT image analysis

The nuclear staining was used to align the images of the same areas between the hybridizations, using Zen2.5 (Carl Zeiss Microscopy GmbH). The images were analyzed as 16bit *.tiff files, without compression or scaling. Images were tiled using a custom script in Fiji (Preibisch et al., 2009; Schneider et al., 2012). The signal-dots were counted using CellProfiler 4.13 (McQuin et al., 2018), Fiji (Preibisch *et al*., 2009; Schneider *et al*., 2012) and R-RStudio (Allaire, 2012; Peterson et al., 2015; Team, 2013; Wickham, 2009; Wickham and Wickham, 2016) custom scripts (https://github.com/AlexSount/SCRINSHOT-HDCA). The identified signal-dot coordinates were used to project the signals on DAPI images, using TisUUmaps (Solorzano *et al*., 2020).

For the quantitative analysis of SCRINSHOT datasets of all analyzed time point (6, 8.5, 11.5 and 14 pcw) nuclei images were segmented into hexagonal bins of 7µm radius. Only bins with a clear proximal epithelial component, SOX2 dots>5) were further processed. To maintain NE-related bins, we used the analyzed genes that were specifically expressed in NE-cells according scRNA-seq (*ARX, NKX2-2, GHRL, ACSL1, CALCA, GRP, RFX6, CFC1* and PCSK1). Bins with a presence of at least 7 signals of any of the mentioned genes were kept in the analysis. We also kept bins containing more than 4 ASCL1 dots, which was found to be expressed by NE-progenitors.

We performed Leiden clustering with 0.3 resolution in individual samples and represented those clusters using UMAP-plots. We further assessed the correlation in expression between the different neuroendocrine genes kept in the analysis in every individual sample, including *ASCL1* and representing the correlations as heatmap. Finally, the suggested clusters were annotated based on the combination of different NE-markers, according to the scRNA-Seq data and mapped back in the tissue. Temporal dynamics were estimated by calculating the relative frequency of each NE-identity in each time.

### Exploration of the zonation patterns in the developing lung using ISS

To calculate the relative position of distinct cell-types in the proximal-distal and radial axis, analyzed tissues with HybISS were segmented into bins (radius: 20µm). Only bins with more than 3 detected EPCAM mRNAs were considered to be airway-related. We calculated the distance of each bin in the tissue to the closest identified airway-related bin, defining the first axis explored (radial axis considering the airway as the center). Cells with a radial distance higher than 140µm were excluded from the analysis. To define the second axis, we explored the diversity within airway-related bins and, by UMAP-dimension reduction, we identified that the first dimension recapitulated the proximal-distal typical patterning, based on the expression of know markers. We used that value as pseudotime to assign a proximal-distal value to each of the detected bins. These values served as the second axis of the analysis, considering the proximal-distal value of the closest epithelial bin as the proximal-distal value of the analyzed mesenchymal cells. The distribution of the cells analyzed was represented using KDE-based heat maps.

### Exploration of the zonation patterns in the developing lung using ST

To explore the zonation of mesenchymal populations present in the developing lung with ST datasets we analyzed sections from 8.5 pcw. We identified ST spots containing airways by looking at the expression top 10 differentially expressed epithelial markers (Suppl. Fig. 1G). Cells containing more than 8 UMIs were considered as airway-related ST spot. To define the radial axis, each ST spot was given a value depending on its distance from its closer airway-related ST spot. The proximal-distal axis was calculated based on the compared relative expression levels of known proximal (SOX2, SCGB3A2) and distal (ETV5, TPPP3) epithelial markers. Based on the relative expression of proximal and distal markers, every epithelial ST spot was given a value between −1 (proximal) and 1 (distal). ST spots that were not airway-related were given the proximal-distal score of their closest airway-related ST spot. After rounding the proximal-distal scores of every ST spot, the frequency of every cluster detected using Stereoscope was then computed by averaging ST spots with the same proximal-distal and radial coordinates.

### Immunofluorescence

Tissue sections were prepared, using the same protocol as SCRINSHOT (Sountoulidis *et al*., 2020). Fresh frozen material was fixed with 4% PFA for 10 minutes, at room temperature and slides were washed 3 times for 5 min with PBS 1X (pH 7.4) to remove the fixative. We incubated the sections with 5% donkey serum (Jackson ImmunoResearch, 017-000-121) in PBS 1X (pH 7.4) with 0.1% Triton X100 (blocking buffer) for 1 hour at room temperature (RT) and then they were incubated with primary antibodies in blocking buffer O/N at 4°C. Slides were washed with PBS 1X (pH 7.4), 3 times for 5 min and incubated with secondary antibodies in 2% donkey serum in PBS 1X (pH 7.4) with 0.1% Triton X100 for 1 hour at room temperature. After 3 washes with PBS 1X (pH 7.4) for 10 minutes/each, nuclei were counterstained with 0.5µg/ml DAPI (Biolegend, 422801) in PBS 1X (pH 7.4) in 0.1% Triton X100 and slides were mounted with ProLong Diamond Antifade Mountant (Thermo, P36961) and coverslipped. For ACSL1-CGRP-CDH1 staining, blocking solution contained 1% BSA in 1X TBS (Thermo, 37520) with 0.2% Triton X100, nuclei were counterstained with 2.86µM DAPI (Invitrogen, D21490) and the slides were mounted with Fluoromount-G (Invitrogen, 00-4958-02). All incubations were performed in a humidity chamber.

The following primary antibodies were used: anti-PHOX2B goat polyclonal antibody (R&D Systems, AF4940-SP, 2µg/mL), anti-DLL3 rabbit monoclonal antibody (Cell Signaling Technology, 71804, 1:50), anti-NF-M mouse monoclonal antibody (DSHB, 2H3, 2µg/ml), anti-COL13A1 rabbit polyclonal antibody (NovusBio, NBP2-13854, 1:100), Cy3 anti-αSmooth Muscle Actin (ACTA2) mouse monoclonal antibody (Sigma Aldrich, C6198, 1:2000), anti-GHRL rat monoclonal antibody (R&D Systems, MAB8200-SP, 1.25µg/ml), anti-GRP rabbit polyclonal antibody (Bioss, bs-0011R, 1:200), anti-SOX10 goat polyclonal antibody (R&D

Systems, AF2864-SP, 5µg/ml), anti-MASH1 rabbit monoclonal antibody (Abcam, ab240385, 1:100), anti-ISL1 mouse monoclonal antibody (DSHB, 40.2D6, 1.3µg/ml), anti-CGRP mouse monoclonal antibody (Santa Cruz, sc57053, 1:250), anti-ACSL1 rabbit polyclonal antibody (Atlas Antibodies, HPA011964, 1:250) and anti-E-cadherin mouse Alexa Flour 555 (BD Biosciences, 560064, 1:100). For visualization the following secondary antibodies were applied: Alexa Fluor 488 donkey anti-goat (Jackson ImmunoResearch, 705-545-147, 1:400), Cy3 donkey anti-rabbit (Jackson ImmunoResearch, 711-165-152, 1:400), Cy5 donkey anti-mouse (Jackson ImmunoResearch, 715-175-151, 1:400), Cy5 donkey anti-rabbit (Jackson ImmunoResearch, 711-175-152, 1:400), Alexa Fluor 488 donkey anti-rat secondary antibody (Jackson ImmunoResearch, 712-545-153, 1:400), Alexa Fluor 647 goat anti-mouse (Invitrogen, A32728, 1:250) and Alexa Fluor 488 goat anti-rabbit (Invitrogen, A11034, 1:250) antibodies.

Sections treated with anti-PHOX2B goat, anti-DLL3 rabbit, anti-COL13A1 rabbit and Cy3 anti-Actin, α-Smooth Muscle (ACTA2) mouse monoclonal antibodies were incubated in TE-buffer (10mM Tris, 1 mM EDTA pH 9.0) for 30 minutes, at 80°C in a waterbath and cooled on ice for 30 minutes to facilitate antigen retrieval and washed 3 times for 5 min with PBS 1X (pH 7.4), prior incubation with the blocking solution.

### Image acquisition for immunofluorescence

Image acquisition was initially done as in SCRINSHOT, with a 10X lens, allowing the identification of informative regions of interest. For high resolution images, we used a Zeiss LSM800 confocal microscope, equipped with a Plan-Apochromat 40X/1.30 oil lens. Optimal resolution settings were used and images were acquired as optical stacks. For imaging of the ACSL1-CGRP-CDH1 stainings, we used a Leica DMI8 microscope (Leica Microsystems, 11090148013000), with a SOLA light engine light source (Lumencor,16740), equipped with a 40X/ 0.80 HC Fluotar, a Hamamatsu camera (2048 × 2048, 16-bit, C13440-20C-CL-301201) and an automated stage (ITK Hydra XY). For the signal detection, we used the following Chroma filters: QUAD-S filter set: DFTC (DC:425; 505; 575; 660). Imaging was done via the LASX software (Leica Microsystems) and images were analysed with Fiji (Preibisch *et al*., 2009; Schneider *et al*., 2012).

### Browser-based interactive visualization of the scRNA-Seq, spatial and interactome analyses

For the browser-based representation of our data, we used the TissUUmaps tool (Solorzano *et al*., 2020). In the presented version, we have modified TissUUmaps for accelerated GPU-based rendering, enabling real time interactive multi-scale viewing of millions of data points directly via a web browser. Furthermore, we have added functionality so that ST-data and single cell pciSeq data from ISS can be presented as pie-charts for efficient viewing of spatial heterogeneity. TissUUmaps supports FAIR sharing of data by allowing users to select regions of interest and directly download raw data in a flexible *.csv format, enabling further exploration and analysis, of all datasets. We based the interactome browser in the Cell Chat shiny app, described in (Jin *et al*., 2021)

## Supporting information

Differential_expression_analyses

SCRINSHOT_padlock_probes

ISS_padlock_probes

## Acknowledgments

We thank National Genomics Infrastructure for sequencing services, the Karolinska Institutet Developmental Tissue Bank for providing human prenatal tissue and the In Situ Sequencing (ISS) facility, at SciLifeLab for ISS service. This work was supported by grants from the Knut and Alice Wallenberg Foundation (KAW 2018.0172), the Erling Persson Foundation, the Chan Zuckerberg Initiative (SVCF 2017-173964), Cancerfonden (MN: CAN 2018/604) and The Swedish research council (MN: 2019-01238) and grants from Cancerfonden and the Swedish Research Council to C.S.

## Figure Legends

**Supplementary Figure 1.**
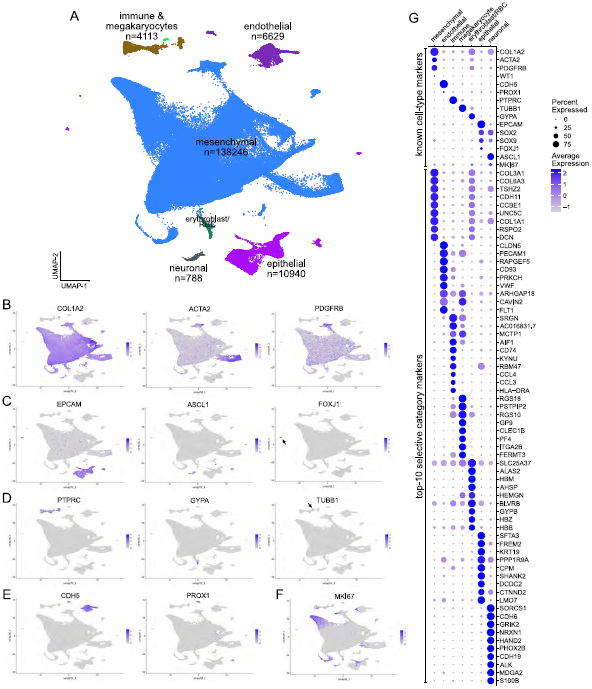
Initial scRNA-Seq analysis suggests 6 main cell classes, with distinct gene-expression profiles. **(A)** Whole dataset UMAP-plot of the 6 main suggested cell-classes. “n”: number of cells/category. **(B-F)** UMAP-plots showing the expression levels of characteristic cell-type marker. (B) *COL1A2*: Mesenchymal (4), *ACTA2*: ASM-pericytes (24), *PDGFRB*: pericytes (25), (C) *EPCAM*: Epithelial (26), *ASCL1*: Neuronal (27, 28) and Neuroendocrine (29), (D) *PTPRC*: Immune (10), *GYPA*: Erythoblast/erythrocyte (30), *TUBB1*: Megakaryocyte (23), (E) *CDH5*: Endothelial (5), *PROX1*: Neuroendocrine (31) and Lymphatic Endothelial (7), (F) *MKI67*: proliferating (8). Expression levels: log_2_(normalized UMI-counts+1) (library size was normalized to 10.000). Blue: high, Gray: zero expression **(G)** Balloon-plot showing the expression of known cell-type markers and the top-10 most selective category markers.

**Supplementary Figure 2.**
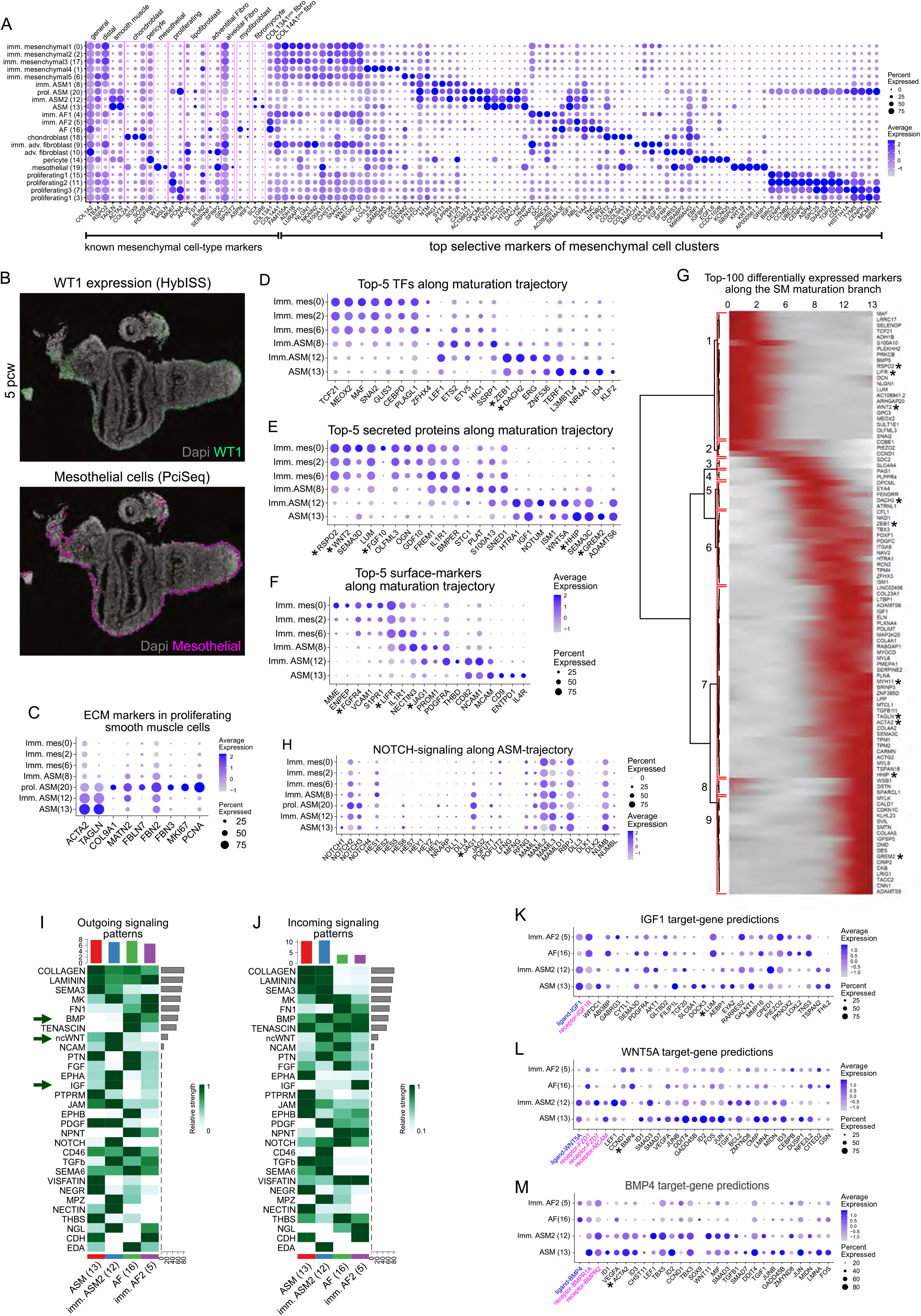
Analysis of mesenchymal cell heterogeneity. **(A)** Balloon-plot of known mesenchymal cell markers (*COL1A2-COL14A1*). General: *COL1A2* (Travaglini *et al*., 2020), *TBX4* (Kumar *et al*., 2014), Distal: *RSPO2* (Hein *et al*., 2021), Smooth muscle: *TAGLN, ACTA2* (Travaglini *et al*., 2020), Chondroblast: *COL2A1, SOX9, SOX6* (Liu and Lefebvre, 2015; Zhao et al., 1997), Pericyte: *PDGFRB* (Greif *et al*., 2012), Mesothelial: *WT1* (Cano et al., 2013), *MSLN* (Rinkevich et al., 2012), Proliferating: *MKI67* (Schonk et al., 1989), *PCNA* (Bologna-Molina *et al*., 2013), Lipofibroblast: *APOE, FST, PLIN2* (Travaglini *et al*., 2020), Adventitial fibroblast: *SERPINF1, SFRP2* (Travaglini *et al*., 2020), Alveolar fibroblast: *GPC3, SPINT2* (Travaglini *et al*., 2020), Myofibroblast: *ASPN, WIF1* (Travaglini *et al*., 2020), Fibromyocyte: *SCX, LGR6* (Travaglini *et al*., 2020), COL13A1^pos^ fibroblast: *COL13A1* (Raredon *et al*., 2019) and COL14A1^pos^ fibroblast: *COL14A1* (Raredon *et al*., 2019). The remaining genes correspond to the top-5, most selective genes for each cluster. **(B)** Representative image of *WT1* mRNA expression (top) in lung periphery, where mesothelium is localized (35, 38). Prediction of mesothelial-cell spatial distribution (bottom) by PciSeq, using HybISS data. **(C)** Balloon-plot of the ASM markers *ACTA2* and *TAGLN*, the ECM-proteins *COL9A1, MATN2, FBLN7, FBN2* and *FBN3* and the proliferation markers *MKI67* and *PCNA*, showing increased ECM-protein expression by proliferating ASM. **(D-F)** Balloon-plots of the top-5 transcription factors (TFs) (D), secreted (E) and surface proteins (F), identified by differential expression analysis of the indicated clusters (Fig. 2A). In all cases, the top-10 genes (based on average log2 fold-change) were sorted according to the percent of positive cells and the top-5 markers were plotted. Gene order: cluster order. **(G)** Heatmap of the top-100 differentially expressed genes along the maturation trajectory, based on tradeSeq [4]. Numbers: stable gene-modules. **(H)** Balloon-plot of NOTCH-signaling genes expression in the ASM-differentiation trajectory clusters. **(I-J)** Heatmaps of the incoming (I) and outgoing (J) signaling patterns of the analyzed ASM and AFs, as indicated by CellChat. Bars represent the outgoing/incoming overall potential on each cluster (top) and pathway (right). **(K-M)** Balloon-plots of the top-20 predicted *IGF1* (K), *WNT5A* (L) and *BMP4* (M) -target genes, expressed in the ASM and AF clusters and predicted by NicheNet [2]. Ligands: blue. Receptors: magenta. Balloon size: percent of positive cells. Color intensity: scaled expression. inhibitors (Ouadah et al., 2019). Brackets highlight JAG1 and DLL3.

**Supplementary Figure 3.**
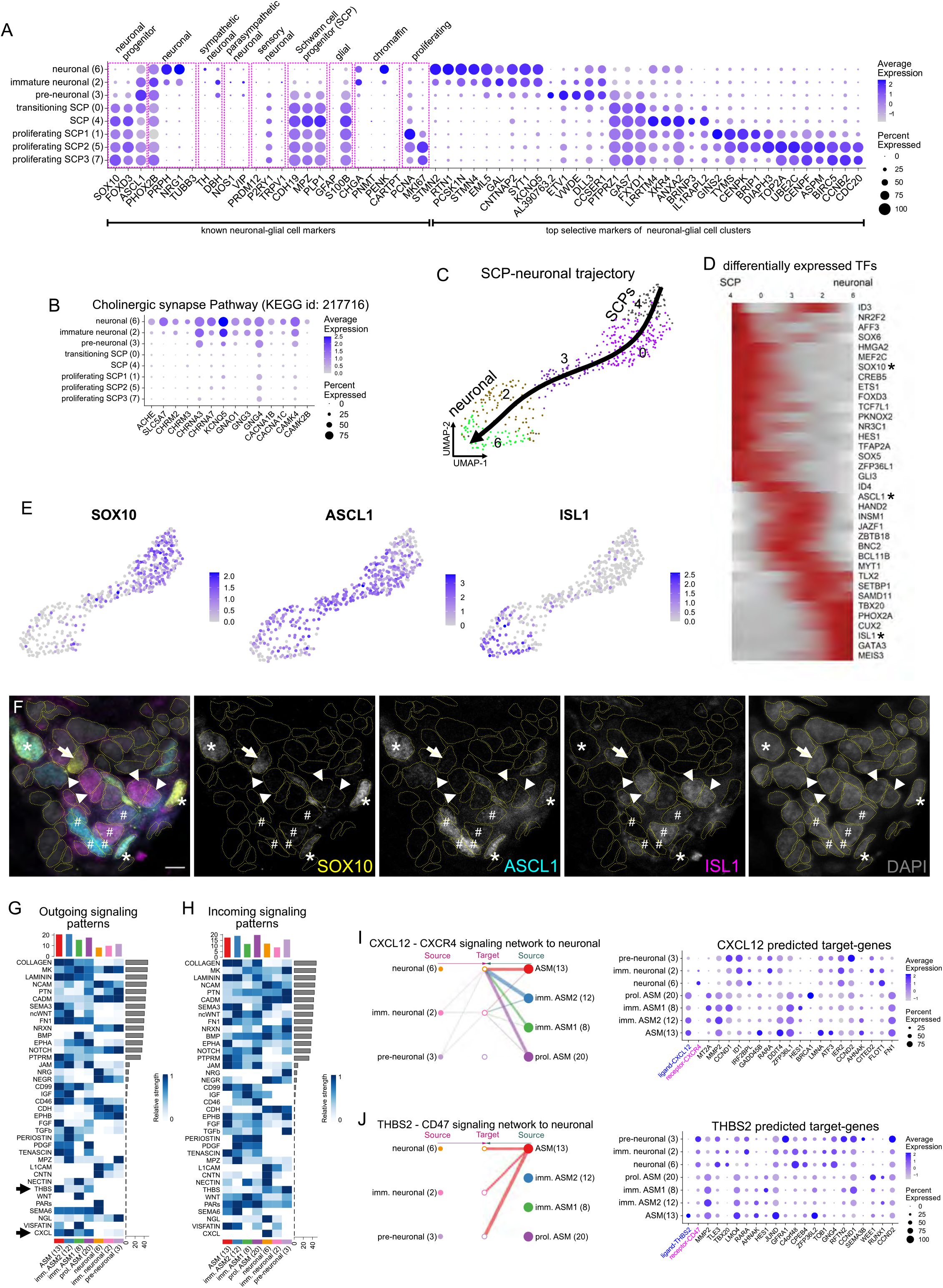
Signaling pathways involved in neuronal cell communications. **(A)** Balloon-plot of known neuronal-glial cell markers (*SOX10-MKI67*). Progenitor: *SOX10* (Kim et al., 2003), *FOXD3* (Simoes-Costa et al., 2012), *ASCL1* (Dyachuk *et al*., 2014), Neuronal: *PHOX2B* (Bielle et al., 2012), *PRPH* (Leung et al., 2004), *NRG1* (Birchmeier and Nave, 2008), *TUBB3* (Sullivan and Cleveland, 1986), Sympathetic neurons: *DBH, TH* (Ernsberger et al., 2000), Parasympathetic neurons: *NOS1, VIP* (Alm et al., 1995), Sensory neurons: *PRDM12, P2RY1, TRPV1* (Chang *et al*., 2015; Kupari et al., 2019), Schwann Cell Progenitors (SCPs): *CDH19, MPZ, PLP1* (Kim et al., 2017), Glial cells: *GFAP, S100B* (Jessen and Mirsky, 2005; 2019), Chromaffin cells: *PNMT, PENK, CARTPT* (Kameneva *et al*., 2021) and Proliferating cells: *MKI67* (Schonk *et al*., 1989), *PCNA* (Bologna-Molina *et al*., 2013). The remaining genes correspond to the top-5, most selective genes for each cluster. **(B)** Balloon-plot of the detected cholinergic-synapse pathway genes (KEGG id: 217716). **(C)** Pseudotime analysis of the SCP-neuronal trajectory on a UMAP-plot, with Slingshot. **(D)** Heatmap of differentially expressed transcription factors (TFs) along the SCP-neuronal trajectory, according to tradeSeq [4]. Stars: analyzed genes in “E-F”. **(E)** UMAP-plots of *SOX10, ASCL1* and *ISL1*. Expression levels: log_2_(normalized UMI-counts+1) (library size was normalized to 10.000). Blue: high. Gray: zero. **(F)** Confocal-microscopy image of an 8.5 pcw ganglion, showing SOX10, ASCL1 and ISL1 expression, detected with immunofluorescence. Dashed outlines: manually segmented nuclei. SOX10^pos^ SCPs (arrows), SOX10^pos^ ASCL1^pos^ transitioning SCPs (asterisks), ASCL1^pos^ SOX10^neg^ immature neurons (hashes), ISL1^pos^ ASCL1^neg^ mature neurons (arrowheads). Scale-bar: 5µm. **(G-H)** Heatmaps of the significant outgoing (G) and incoming (H) signaling patterns between neuronal and airway smooth muscle (ASM) clusters, indicated by CellChat. Bars show the outgoing/incoming overall potential on each cluster (top) and pathway (right). Arrows: further analyzed communication patterns. **(I)** (left) CellChat hierarchical plot of the *CXCL12*-*CXRC4* communication pattern. (right) Balloon-plot of the top-20 *CXCL12*-target genes, predicted by NicheNet. **(J)** as in “I” for *THBS2*-*CD47* pair. Ligands: blue, Receptors: magenta. Balloon size: percent of positive cells. Color intensity: scaled expression.

**Supplementary Figure 4.**
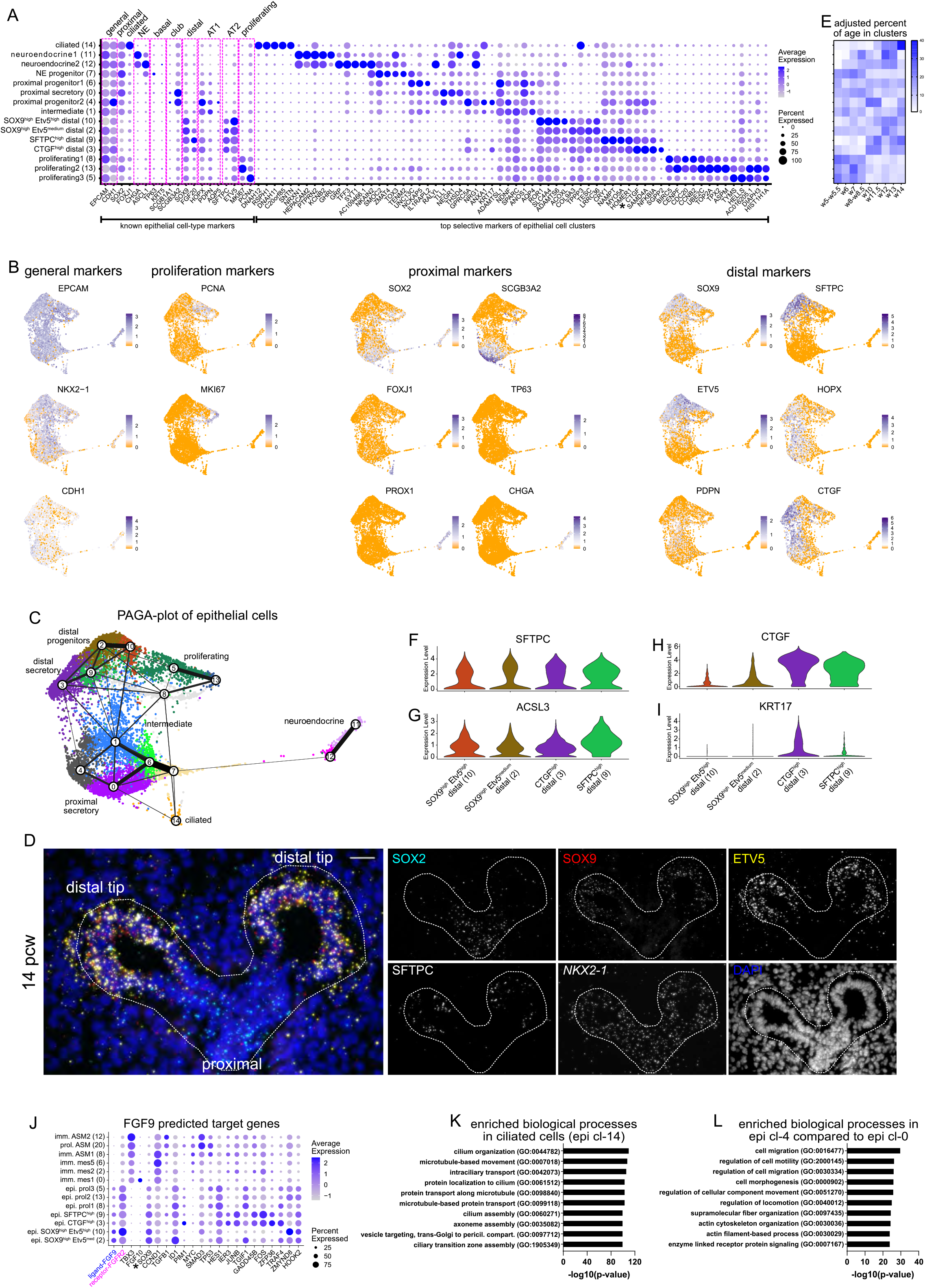
Analysis of epithelial cell heterogeneity. **(A)** Balloon-plot of known epithelial markers in the clusters of Fig. 4A. General: *EPCAM, CDH1*, Proximal: *SOX2* (Nikolic *et al*., 2017), Ciliated: *FOXJ1* (Gomperts et al., 2004), Neuroendocrine: *CHGA, ASCL1* (McGovern et al., 2010), Basal: *TP63, KRT5* (Evans et al., 2001), Club cells: *SCGB1A1, SCGB3A2* (Reynolds *et al*., 2002), Distal: *SOX9* (Nikolic *et al*., 2017), *FGF20*, Alveolar Type 1 (AT1): *HOPX, PDPN, AQP5* (Nikolic *et al*., 2017), AT2: *SFPTC, ETV5* (Zhang et al., 2017) and Proliferating cells: *MKI67* (Schonk *et al*., 1989), *PCNA* (Bologna-Molina *et al*., 2013) markers in addition to the expression of the top-5 differentially expressed genes of each cluster. Balloon size: percent of positive cells. Color intensity: scaled expression. **(B)** UMAP-plots of epithelial cells, showing the expression levels of general markers: *EPCAM, NKX2-1, CDH1*, for proliferation: *PCNA* and *MKI67*, for proximal epithelium: *SOX2, FOXJ1, PROX1, SCGB3A2, TP63, CHGA* and for distal: *SOX9, ETV5, PDPN, SFTPC, HOPX* and *CTGF*. Expression levels: log_2_(normalized UMI-counts+1) (library size was normalized to 10.000). Blue: high, orange: zero. **(C)** PAGA-plot of the analyzed epithelial cells, superimposed on their UMAP-plot. Line thickness shows the probability of cluster-connections. **(D)** Region of interest showing a 14 pcw distal epithelial airway, analyzed with SCRINSHOT. *SOX2* (cyan), *SOX9* (red), *ETV5* (yellow), *SFTPC* (gray), *NKX2-1* (shown only as individual channels) and DAPI (blue). Dotted outline indicates the airway bonders. **(E)** Heatmap of percentages of age abundances in epithelial clusters. To avoid bias, cell numbers were firstly normalized according to dataset size per age. **(F-I)** Violin plots of *SFTPC* (F), *ACSL3* (G), *CTGF* (H) and *KRT17* (I) expression levels in the four distal epithelial clusters. Expression levels: log_2_(normalized UMI-counts+1) (library size was normalized to 10.000). **(J)** Balloon-plot of the top-20 predicted target genes (by NicheNet) of *FGF9*. Balloon size: percent of positive cells. Color intensity: scaled expression. **(K)** Bar-plot of the most enriched biological processes p-values in ciliated cells (epi cl-14). **(L)** Same as in “k” for the proximal progenitor cells (epi cl-4) compared to the proximal secretory (epi cl-0).

**Supplementary Figure 5.**
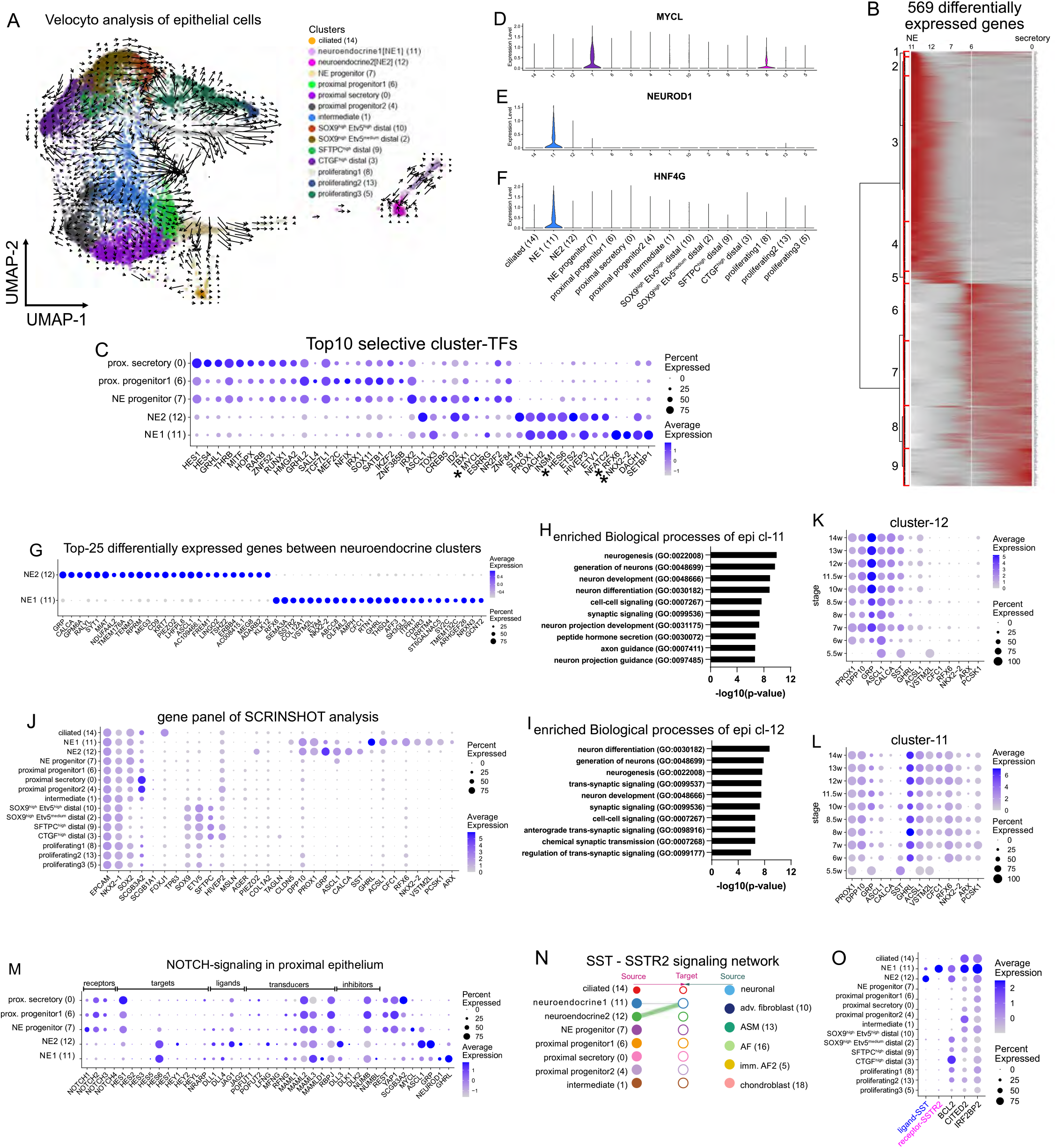
Exploring the diversity within the proximal airway neighborhood. **(A)** RNA velocity analysis of epithelial cells, on the UMAP-plot. Colors as in Fig. 4A and arrow direction indicates the future state of the cells. **(B)** Heatmap of the 569 differentially expressed genes (FDR<0.0001 and FcMedian>1) along the proximal secretory and neuroendocrine (NE) trajectories, based on tradeSeq. The dendrogram (left) indicates 9 stable gene-modules. **(C)** Balloon-plot of the top-10 selective transcription factors (TFs) in the proximal epithelial secretory and NE-clusters. **(D-F)** Violin-plots of the transcription factors *MYCL* (D), *NEUROD1* (E) and *HNF4G* (F), in all epithelial clusters. Expression levels: log_2_(normalized UMI-counts+1) (library size was normalized to 10.000). **(G)** Balloon-plot of the top-25 selective markers for each NE-cluster compared to the other. Balloon size: percent of positive cells. Color intensity: scaled expression. **(H)** Bar-plot of the top-10 enriched biological process p-values in epi cl-11 compared to epi cl-12, using its statistically-significant (adjusted p-value <0.001) upregulated genes. **(I)** as in “H” for epi cl-12, compared to Sam2 as in”L” for epi cl-11. **(J)** Balloon-plot of the 31 analyzed genes with SCRINSHOT. **(K)** Balloon-plot of the NE-gene expression changes over time, in epi cl-12, for the common NE-markers: *PROX1, DPP10*, epi cl-12 markers: *GRP, ASCL1, SST* and *CALCA* and eight epi cl-11 markers *GHRL*-*PCSK1*. **(L)** Same as in “L” for epi cl-11. Balloon size: percent of positive cells. Color intensity: log_2_(normalized UMI-counts+1) (library size was normalized to 10.000). Blue: high. Gray: zero. **(M)** Balloon-plot of NOTCH-signaling components (Ouadah *et al*., 2019), in addition to the neuronal gene inhibitor *REST* (Lim *et al*., 2017), the TF *YAP1*, the secretory marker *SCGB3A2*, and the NE-markers *MYCL, ASCL1, GRP, NEUROD1* and *GHRL*. Balloon size: percent of positive cells. Color intensity: scaled expression **(N)** CellChat hierarchical plot of SST-SSTR2 communication pattern between the two NE-cell states. **(O)** Balloon-plot of *BCL2, CITED* and *IRF2BP2* expression levels that are included in the top-100 NicheNet predicted *SST* target-genes. Ligand: blue, Receptor: magenta. Balloon size: percent of positive cells. Color intensity: scaled expression.

**Supplementary Figure 6.**
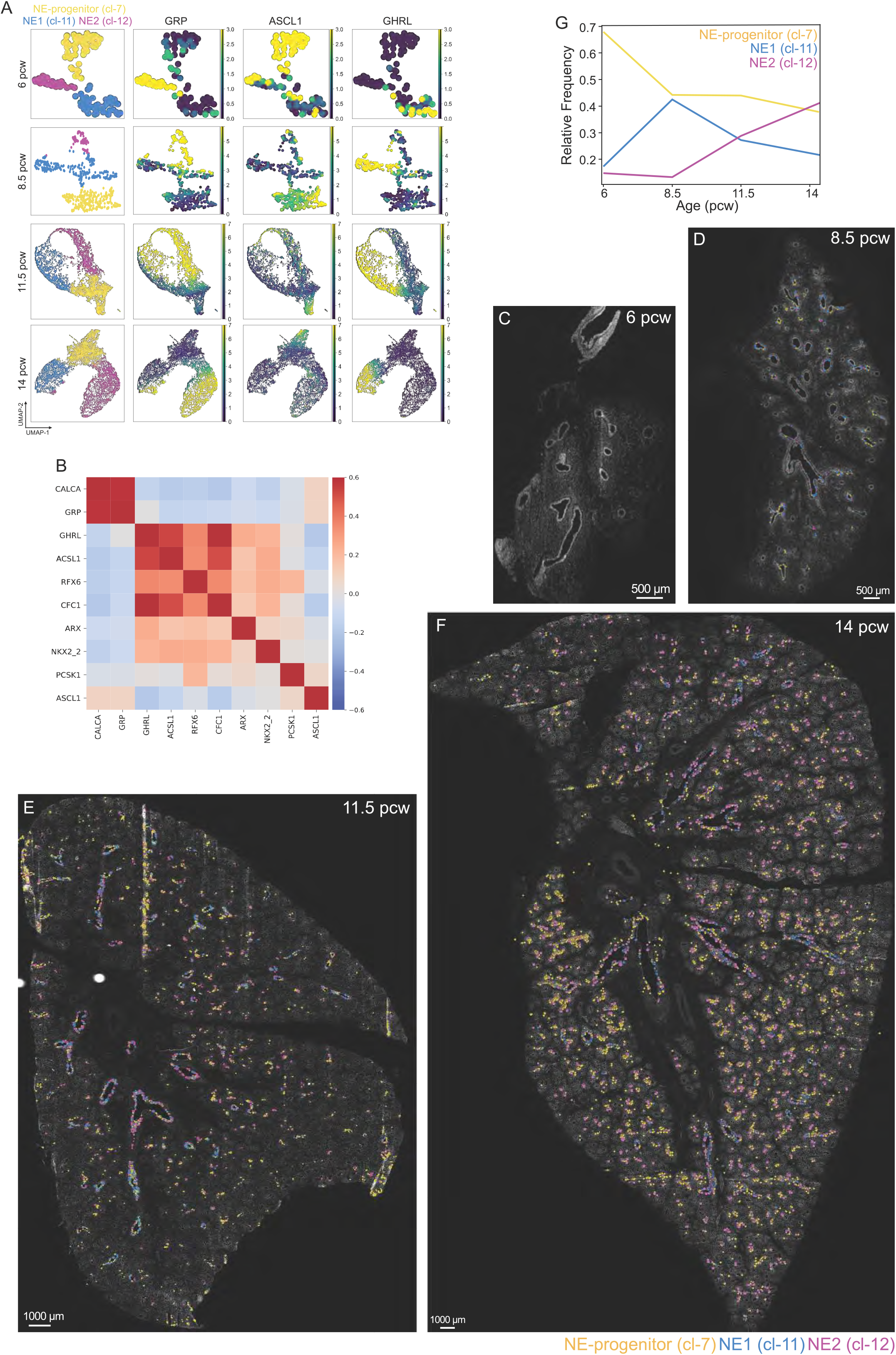
NE-cell distributions in time and space. **(A)** UMAP-plots of neuroendocrine-related bins (Methods) and their progenitors in the analyzed time points with SCRINSHOT, showing the suggested clusters (first column) and the detected mRNAs for *GRP* (second column), *ASCL1* (third column) and *GHRL* (forth column). **(B)** Heatmap of the correlation in expression of the different NE-associated markers included in the SCRINSHOT analysis of a pcw 14 section (shown in G). **(C-F)** Maps of indicated NE-populations according to SCRINSHOT analysis in 6 (C), 8.5 (D), 11 (E) and 14 (F) pcw lung sections. DAPI: gray, NE-progenitor (cl-7): yellow, NE1 (cl-11): cyan, NE2 (cl-12): magenta. **(G)** Line plot of the changes in relative frequency of the three main neuroendocrine-related clusters, across different age sections, analyzed with SCRINSHOT.

**Supplementary figure 7.**
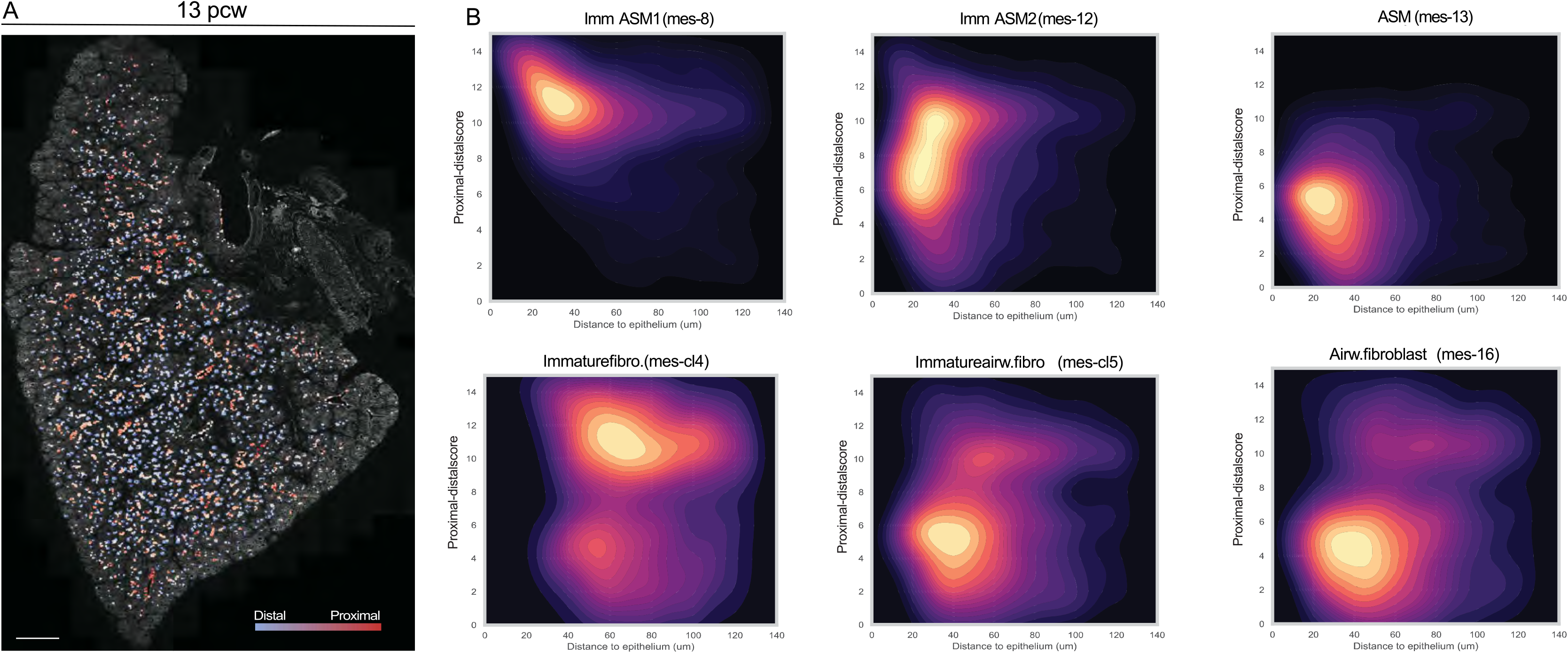
A. Proximal-distal axis score of the epithelium of a 13 pcw lung section, analyzed by HybISS. DAPI: gray, proximal: red, distal: blue. Scale-bar: 1000 µm **(B)** Density maps of ASM and AF clusters, showing their distribution along proximal-distal axis (y-axis) and their distance from the epithelium (x-axis). Brighter colors represent higher relative frequency.

## Supplementary Note

### Immune cells colonize the lung during the first trimester of gestation, showing signs of functional competence

The adult human lung contains a large repertoire of immune cells (Travaglini et al., 2020) that play crucial roles in host defense mechanisms (Lloyd and Marsland, 2017). During development, hematopoiesis takes place in distinct places (yolk sac, aorta gonad mesonephros (AGM), liver and finally bone marrow). In the first trimester of gestation, liver is the main source of immune cells that seed various organs, including lung, to produce their resident immune cells (Popescu et al., 2019).

Our dataset contains 4113 PTPRC^pos^ (Hermiston et al., 2003) cells of IL1B^pos^ myeloid (cl-17) (Di Paolo and Shayakhmetov, 2016) and CD247^pos^ lymphoid (cl-22) immune cell-types, in addition to TUBB1^pos^ megakaryocytes (cl-28) (Freson et al., 2005) (Suppl. Fig. 1A). Their re-clustering suggested 22 clusters (Add. Fig. 1A), which were annotated to distinct immune cell-types and their corresponding immature states, based on the expression of known markers (Add. Fig. 1B). In summary, we identified myeloid macrophages, monocytes, neutrophils, mast/basophis and dendritic cells (DCs) and lymphoid B-cells, innate lymphoid cells (ILCs), and natural killer (NK) cells, in addition to megakaryocytes. Myeloid cells are more abundant in early stages of development and lymphoid appear later, after 10 pcw (Add. Fig. 1C, D).

Macrophages is the most numerous immune cell-type, that includes two immature (−0 and - 22) and one mature cluster (cl-16), that expresses highly TNF, suggesting that embryonic lung contains functionally competent macrophages (Parameswaran and Patial, 2010). Examination of known myeloid-cell effector proteins, such as the cytokines IL1A, IL1B (Di Paolo and Shayakhmetov, 2016), IL23A (Pirhonen et al., 2002) and the chemokines CCL4 and CXCL2 (Mantovani et al., 2004) provided additional evidence that not only macrophages but also monocytes (immune cl-14) can be operative (Add. Fig. 1B, E). The myeloid lineage also included: (i) the MPO^pos^ (Klebanoff et al., 2013) neutrophils, with a small percentage of them being ELANE^pos^ (Takahashi et al., 1988), suggesting that they are still immature (Fig. 5 B), (ii) MKI67^pos^ PCNA^pos^ proliferating myeloid cells (immune cl-8) (Add. Fig. 1B) and (iii) three types of dendritic cells that all express HLA-DQA1 (Travaglini *et al*., 2020) but only cl-5 expresses CCR7, suggesting that it contains migrating dendritic cells (Riol-Blanco et al., 2005) and only cl-19 expresses CLEC9A, a conventional dendritic cell marker (Yamasaki, 2016).

The lymphoid lineage contained five distinct clusters (Add. Fig. 1A). The NKG7^pos^ NCR1^pos^ KLRD1^pos^ NK-cells is the most abundant cell-type (immune cl-1) (Fig. 5 B). Its positivity for previously reported lymphoid-cell effector proteins, like IFNG (Guo et al., 2018), GNLY and the granzymes A and M (Dotiwala and Lieberman, 2019) indicates that embryonic lung NK-cells are functionally potent (Add. Fig. 1F). The immune cl-13 corresponds to GATA3^pos^ IL1RL1^pos^ innate lymphoid cells 2 (ILC2) and the cluster-21 to the IL1R1^pos^ IL23R^pos^ ILC3. While both are KIT^pos^ (Meininger et al., 2020) (Add. Fig. 1B), only ILC2 produces the cytokine IL13, as previously reported (Nagasawa et al., 2019) (arrow in Add. Fig. 1F).

**Additional Figure 1.**
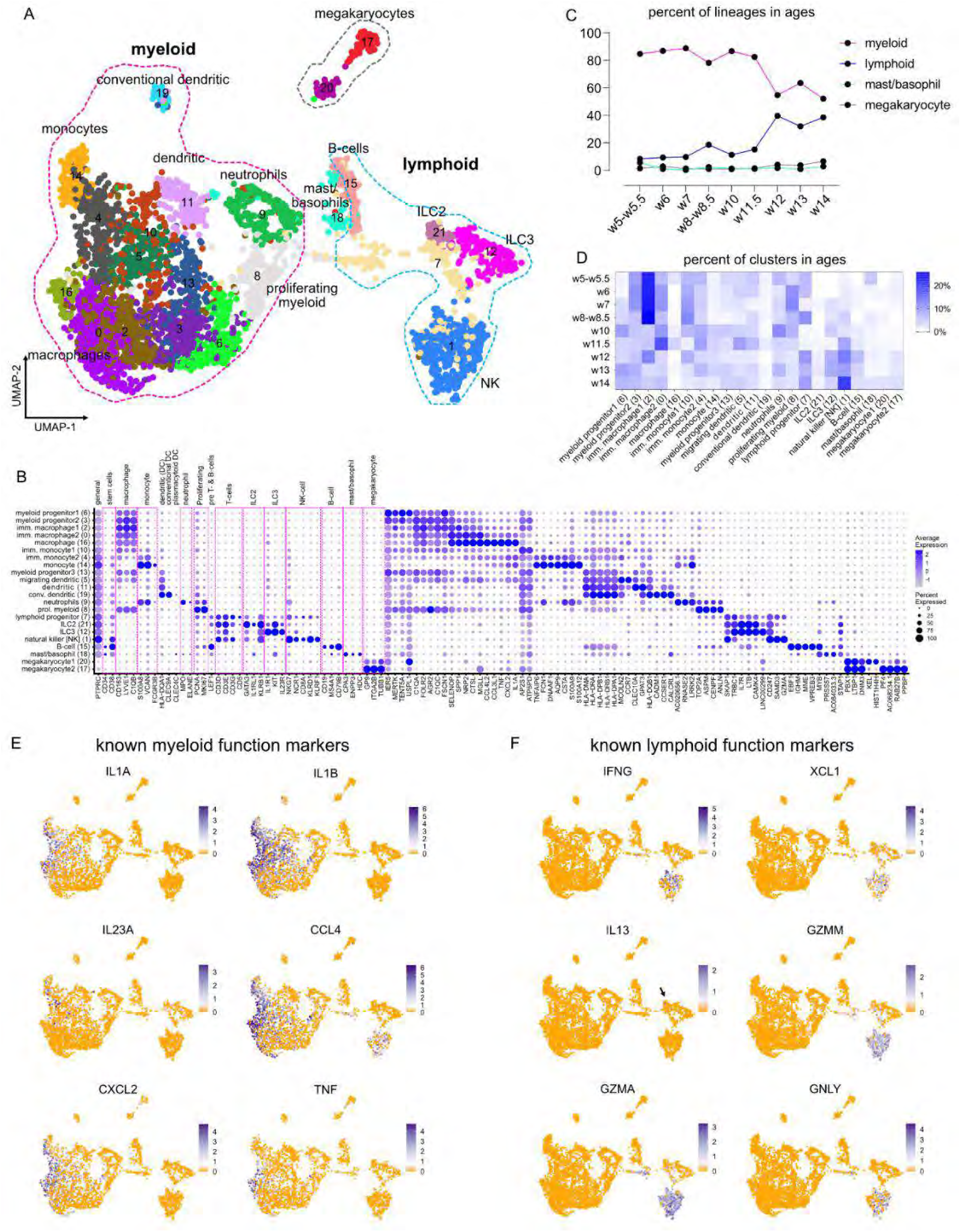
Immune cell characterization suggests thar all main cell-types are found in the embryonic lung, being functionally competent. **(A)** UMAP-plot of the 4113 analyzed immune cells of the whole dataset. Different colors indicate the 22 suggested clusters. Dotted outlines show the myeloid (magenta), lymphoid (cyan) lineages and megakaryocytes (gray). **(B)** Balloon-plot of known immune cell markers (PTPRC-TUBB1). General: PTPRC (Hermiston *et al*., 2003), Stem cells: CD34, CD38 (Popescu *et al*., 2019), Macrophages: CD163, LYVE1 (Chakarov et al., 2019), C1QB (Travaglini *et al*., 2020), Monocytes: S100A8, VCAN (Travaglini *et al*., 2020), FCGR3B (Bertrand et al., 2004), Dendritic cells: HLA-DQA1 (general), CLEC4C (plasmacytoid) (Travaglini *et al*., 2020), CLEC9A (myeloid) (Canton et al., 2021), Neutrophils: MPO, ELANE (Popescu *et al*., 2019), Proliferating: MKI67 (Schonk et al., 1989), PCNA (Bologna-Molina et al., 2013), T-cells: CD3D, CD3E, CD3G (Dong et al., 2019; Travaglini *et al*., 2020), CD5 (Domingues et al., 2016), ILC2: GATA3, IL1RL1 (Dahlgren et al., 2019), KLRB1 (CD161) (Bal et al., 2020), ILC3: IL1R, IL23R, KIT (Spits et al., 2013), NK-cells: NKG7, NCR1, KLRD1, KLRF1 (Travaglini *et al*., 2020), CD8A (Popescu *et al*., 2019), B-cells: CD19, MS4A1, CD79B (Travaglini *et al*., 2020), Mast/basophils: CPA3, HDC (Dwyer et al., 2016), ENPP3 (Buhring et al., 2004) and GP9 (Bianchi et al., 2016), ITGA2B (Travaglini *et al*., 2020), TUBB1 (Freson *et al*., 2005). The remaining genes correspond to the top-5, most selective genes for each cluster. Balloon size: percent of positive cells. Color intensity: scaled expression. **(C)** Line plot of lineage abundancies (%) in the analyzed ages. **(D)** Heatmap of cluster abundancies (%) in the analyzed ages. **(E-F)** UMAP-plots of gene expression levels of known myeloid (E) and lymphoid (F) function genes. Expression levels correspond to log_2_(normalized UMI-counts+1) (library size was normalized to 10.000). Blue: high, Orange: zero.

The lymphoid cl-7 is positive for the general T-cell markers CD3D, CD3G, CD3G (Dong *et al*., 2019) and presumably corresponds to immature lymphoid cells and the cl-15 contains CD19^pos^ CD79B^pos^ MS4A1^pos^ B-cells (Add. Fig. 1B). The cl-18 expresses basophil and Mast-cell markers like CPA3, HDC (Dwyer *et al*., 2016) and ENPP3 (Buhring *et al*., 2004) but the low percent of positive cells in the cluster suggests that they are still immature (Add. Fig. 1B). Finally, we detected two populations of GP9^pos^ (Bianchi *et al*., 2016) megakaryocytes, which might correspond to immature (immune cl-20) and mature (immune cl-17) cell-states, based on the expression levels of the examined differentiation markers (Add. Fig. 1B).

In summary, our analysis provides a detailed description of the immune cell-types of the developing lung during the first trimester of gestation. We cannot answer whether these cells derive from the circulation or consist already-established, resident cells of the organ, like alveolar macrophages (Hussell and Bell, 2014) and ILCs (Dahlgren *et al*., 2019) but we confirmed that the major immune cell-types can access the developing organ, earlier than previously believed (de Kleer et al., 2016; Michaelsson et al., 2006) and be functional, as indicated by the expression of distinct effector proteins, at least, at the mRNA level.

### The main lung endothelial cell-types appear during the early stages of lung development

Focusing on the mesoderm-derived cell-types, we individually analyzed the endothelial cells (Suppl. Fig. 1A). We annotated the suggested clusters to arterial, venous, capillary, lymphatic and bronchial endothelial cells (Add. Fig. 2A), based on already known markers of the adult human lung endothelium (Schupp et al., 2021; Travaglini *et al*., 2020) (Add. Fig. 2B, C) and we showed that these cell-types are already formed during the initial stages of lung development. The positivity of endothelial cluster-9 (endo cl-9) for the CXCL12, IGFBP3, GJA5 and DKK2 indicates that they correspond to arterial cells. The endo cl-13 corresponds to the CPE^pos^ CDH11^pos^ venous endothelium (Schupp *et al*., 2021). The endo cl-2 contains IL7R^pos^ cells, which also express the kinases SGK1 and NTRK2 (Schupp *et al*., 2021; Zarrinpashneh et al., 2013), suggesting that they are capillary endothelial cells. The positivity of endo cl-7 for the PROX1 (Wigle et al., 2002; Wigle and Oliver, 1999) and CCL21 (Travaglini *et al*., 2020) indicates that it contains lymphatic endothelial cells. HBEGF expression was found elevated in endo cl-8, suggesting that it corresponds to bronchial vascular endothelium (Travaglini *et al*., 2020).

Interestingly, the bronchial venous endothelial marker COL15A1 (Schupp *et al*., 2021) was not expressed in that cluster but in immature arterial cl-6, suggesting of differences between embryonic and adult lung endothelium (Add. Fig. 2B). Three of the remaining clusters (−3, −10 and −14) were annotated to proliferating cells, as indicated by their MKI67 (Schonk *et al*., 1989), PCNA (Bologna-Molina *et al*., 2013) positivity (Add. Fig. 2B, C). The rest of the clusters were annotated to immature cell-states, that expressed low levels of differentiation markers, like the CPE^low^ immature venous cl-4 and they were highly positive for the proto-oncogene KIT, confirming previous experimental data about its expression by immature endothelial progenitors of the human embryonic lung (Suzuki et al., 2014). The endo cl-5 showed the highest KIT expression levels, suggesting a type of progenitor (Add. Fig. 2C). The fact that both mature and immature endothelial clusters contain cells from all analyzed stages suggests that lung angiogenesis is asynchronous, at least in first trimester (Add. Fig. 2D).

### The developmental trajectory of arterial endothelium

The arterial endothelial developmental trajectory was one of the most prominent in our analysis. Here, we reasoned to monitor the physiological development of arterial cells, providing a reference, which can facilitate the identification of aberrant, activated mechanisms in disease conditions like the pulmonary arterial hypertension (PAH) (Evans et al., 2021). The arterial developmental trajectory includes the endothelial clusters −12, −6 and −9 (Add. Fig. 2A). In the analysis, we also included the immature clusters −0 and −1 to gain information about the early events of cell specification and differentiation (Add. Fig. 2E). Using pseudotime analysis, we ordered the cells according to their differentiation state, as indicated by the expression patterns of mature endothelial markers, like the CXCL12, DKK2 and SOX5 (Travaglini *et al*., 2020). Our results suggest five statistically-significant modules of the top-50 genes, that are differentially expressed along the trajectory (Add. Fig. 2F). Module-1 contains genes that are expressed by the immature cells of endo clusters −0, −1 and partially −12. This module contains the capillary marker CA4 (Schupp *et al*., 2021). The adult aerocyte marker HPGD (Gillich et al., 2020; Schupp *et al*., 2021) is also included here and might control the endothelial cell migration and vessel formation (Finetti et al., 2008) by inactivating prostaglandins, as it has previously reported (Add. Fig. 2F) (Cho et al., 2006). Immature cells were positive for the TF LHX6, which anti-correlates with arterial cell-identity and has been previous reported to function as a cell-fate mediator, downstream of various signaling pathways, especially in nervous system (Zhou et al., 2015). The module-2 corresponds to upregulated genes in the immature clusters −12 and −6, including the insulin receptor gene (INSR), that controls endothelial cell sprouting, down-stream of VEGF-signaling (Walker et al., 2021). The other three modules contain genes that are active in the mature cells but their activation is asynchronous (Fig. 3 F).

**Additional Figure 2.**
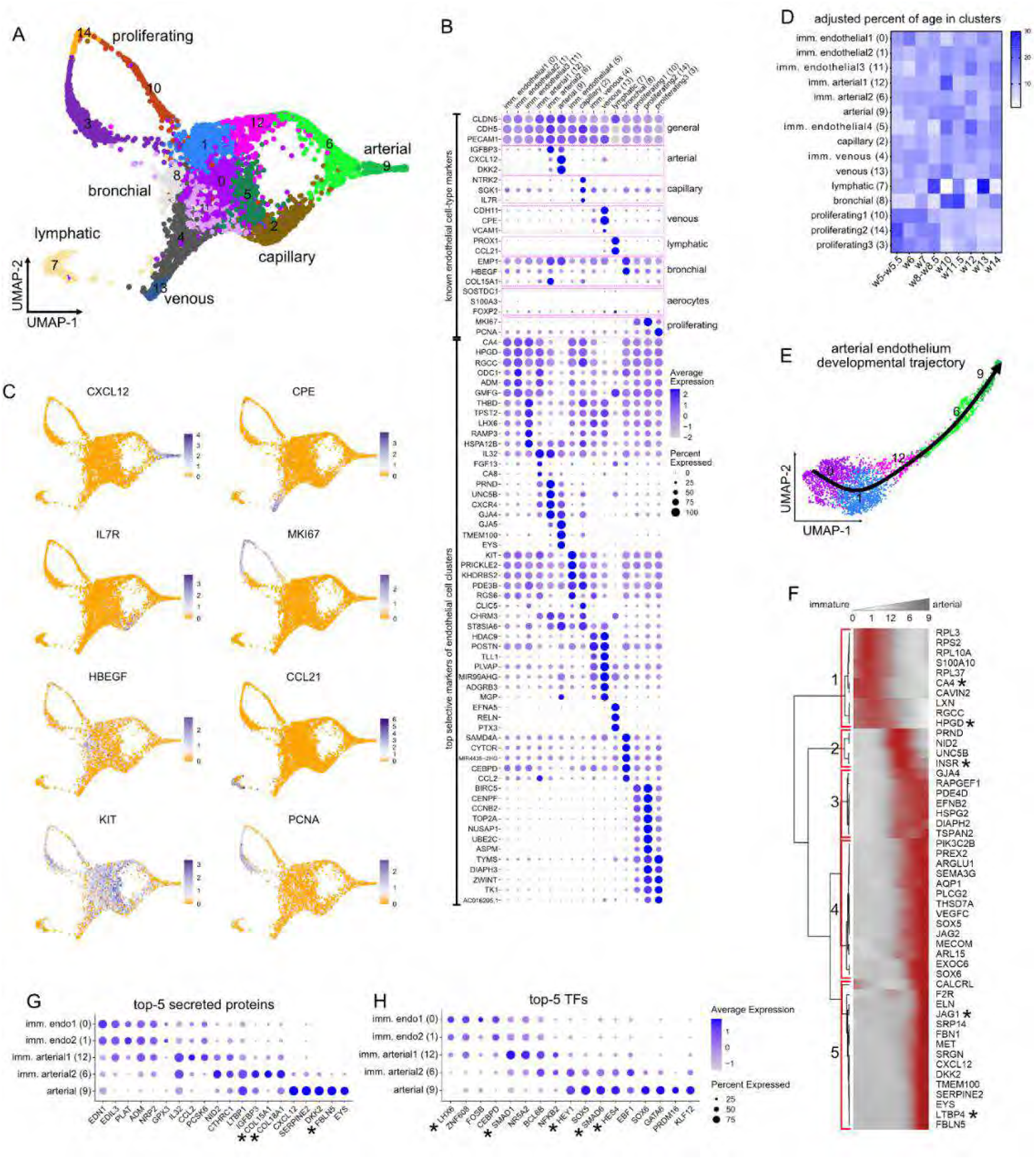
Analysis of endothelial cell heterogeneity. **(A)** UMAP-plot of the 6629 endothelial cells. Different colors indicate the 15 suggested clusters. **(B)** Balloon plot of known endothelial cell markers (CLDN5-PCNA). General: CLDN5, PECAM1 (Travaglini *et al*., 2020), CDH5 (Schupp *et al*., 2021), Arterial: IGFBP3, CXCL12, DKK2 (Travaglini *et al*., 2020), Capillary: NTRK2, SGK1 (Travaglini *et al*., 2020), IL7R (Schupp *et al*., 2021), Venous: CDH11, CPE, VCAM1 (Schupp *et al*., 2021), Lymphatic: PROX1 (Wigle *et al*., 2002; Wigle and Oliver, 1999), CCL21 (Travaglini *et al*., 2020), Bronchial: EMP1, HBEGF (Travaglini *et al*., 2020), COL15A1 (Schupp *et al*., 2021), Aerocytes: SOSTDC1, S100A3 (Travaglini *et al*., 2020), FOXP2 (Schupp *et al*., 2021) and Proliferating: MKI67 (Schonk *et al*., 1989), PCNA (Bologna-Molina *et al*., 2013). The remaining genes correspond to the top-5, most selective genes for each cluster. Balloon size: percent of positive cells. Color intensity: scaled expression. **(C)** UMAP-plots of endothelial cell-type markers, as described in “C”. Expression levels correspond to Iog _2_(normalized UMI-counts+1) (library size was normalized to 10.000). Blue: high, Orange: zero. **(D)** Heatmap of percents of age abundances in endothelial clusters. To avoid bias, cell numbers were firstly normalized according to dataset size per age. **(E)** Pseudotime analysis of the arterial endothelial trajectory, with Slingshot. **(F)** Heatmap of the top-50 differentially expressed genes along the arterial trajectory, according to tradeSeq. The numbers on the left correspond to 5 stable gene-modules (red squares).

Interestingly, the interrogation of the arterial clusters for differentially expressed, secreted proteins showed that the immature arterial cells were positive for the collagen genes, COL15A1 (Schupp *et al*., 2021) and COL18A1, which are down-regulated in the mature state (Add. Fig. 2G). On the contrary, the elastogenesis-related genes FBLN5 and LTBP4 (Add. Fig. 2F, G) (Kumra et al., 2019) were detected in the most mature arterial state of our dataset (endo cl-9).

Focusing on transcription factors, we found that the SMAD1 is upregulated in the immature cl-12. That agrees with its role in positive regulation of many NOTCH-signaling components (e.g., HES4, JAG1 and HEY1) (Add. Fig. 2H) that are necessary for the tip-versus stalk-cell selection in the developing arterial endothelium (Moya et al., 2012). Interestingly, more mature arterial endothelial cells express the inhibitory SMAD6 (Imamura et al., 1997) (Add. Fig. 2H), that functions downstream of NOTCH-signaling to downregulate proliferation and induce junction-associated genes (Ruter et al., 2021).

### Arterial endothelium communicates with pericytes

Based on the spatial co-occurrence of the identified cell-identities, the arterial endothelial cells were found in the same neighborhood as pericytes, adventitial fibroblasts and immature mesenchymal cells of mes cl-4 (Fig. 1H), suggesting cell-cell communications for proper arterial wall development and circulation (Gao and Raj, 2010). Interestingly, the high colocalization between endothelium and pericytes involved only the mature arterial endothelial cells (cl-9), indicating that pericytes cover the vessels only when endothelial cells reach an advance maturation state (Add. Fig. 3A). To better analyze the communication patterns in arterial niche, we focused on the whole arterial endothelial trajectory and the three aforementioned mesenchymal cell identities (Add. Fig. 3B). Here, we confirmed the recently described role of PDGF and EDN pathways in pericyte recruitment and proliferation (Kemp et al., 2020), suggesting conserved mechanisms that are activated during human lung vascular development. Interactome analyses suggested that PDGFB is produced by all analyzed endothelial cells and signals through mainly PDGFRB to pericytes, upregulating the ACTA2 expression (Add. Fig. 3C). EDN1 was expressed by all endothelial cells but was significantly higher in immature endothelial cl-0 and −2. The collagen genes COL1A1, COL1A2 and COL3A1 were included between the top predicted target genes in mesenchymal cells, in addition to PRKCA, that might regulate in the developing artery vasoconstriction (Barman et al., 2004) (Add. Fig. 3D). HGF was the only predicted ligand to be exclusively expressed by pericytes and target the arterial endothelial cells (cl-6, −9), possibly contributing to endothelial cell growth and tube formation, similarly to its role in the rat lung (Seedorf et al., 2016) (Add. Fig. 3E). Our analysis also, suggested that DHH, through PTCH1, induces EGFR expression in all three mesenchymal clusters contributing to their recruitment by endothelial cells for tube formation (Stratman and Davis, 2012) (Add. Fig. 3F).

**Additional Figure 3.**
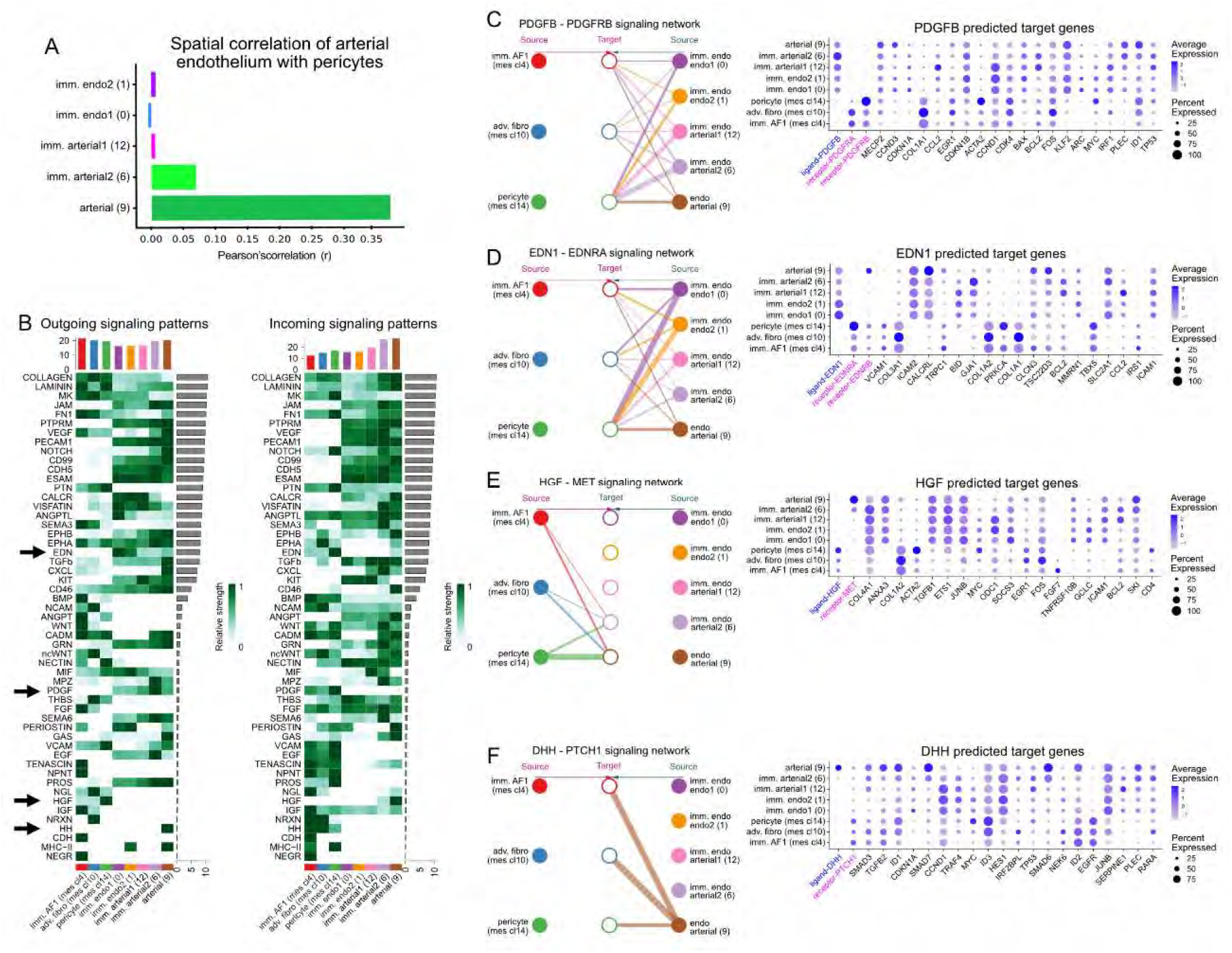
Communication patterns of arterial endothelium. **(A)** Bar-plot of correlation between pericytes (mes cl-14) and the different endothelial clusters of the arterial trajectory, based on ST-data. Colors as in Add. Fig. 2A. **(B)** CellChat (Jin et al., 2021) heatmaps of the incoming and outgoing signaling patterns, between the arterial endothelium and its neighbors, as shown in Fig. 1D. Bars represent the outgoing/incoming overall potential on each cluster (top) and pathway (right). Arrows: further analyzed communication patterns. **(C-F)** (left) CellChat hierarchical plots of ligand-receptor pairs for PDGFB (C), EDN1 (D), HGF (E) and DHH (F). (right) Ballon-plots of the top-20 target genes of the corresponding ligands, predicted by NicheNet (Browaeys et al., 2020). Balloon size: percent of positive cells. Color intensity: scaled expression. Ligand: blue, Receptors: magenta.

### Distinct communication patterns between epithelium and mesenchyme

Focusing on the proximal part of the lung, CellChat analysis (Add. Fig. 4A, B) showed that chondroblasts could communicate with ASM through FGF18-FGFR2 suggesting that smooth muscle has a regulator role in chondrogenesis and might explain the abnormal cartilage ring formation in experimental models of FGF18 ectopic expression, during early mouse lung development (Whitsett et al., 2002) (Add. Fig. 4C). Also, chondroblasts receive signals from AFs, through BMP4, to upregulate the expression of the chondrocyte differentiation regulator SOX9 (Hatakeyama et al., 2004) and of CHST11, an important enzyme for chondroitin sulfate production in chondrocytes (Liu and Lefebvre, 2015) (Add. Fig. 4D).

On the other hand, the epithelium was predicted to receive signals from AFs through the secreted ligand BMP5, up-regulating, for example (i) the Cilia and Flagella Associated Protein 73 (CFAP73) (Travaglini *et al*., 2020) in ciliated cells (epi cl-14) and (ii) the secretory vesicle transport regulator BICDL1 (Schlager et al., 2010) in neuroendocrine cells (epi cl-11 and −12) (Add. Fig. 4E).

Mesenchymal-epithelial interactions are achieved through HGF-signaling. The ligand is produced by the proximal lung fibroblasts, targeting the epithelium, but not ciliated and NE-cells, as indicated by MET (receptor) expression levels. NicheNet predicted that except for its own receptor MET, the HGF induces the expression of CFTR, with a possible function in the transepithelial balance of ions and fluid of the embryonic lung epithelial lumen and provided additional information about the beneficial role of HGF treatment on CFTR maturation and stabilization (Lopes-Pacheco, 2019) (Add. Fig. 4F).

**Additional Figure 4.**
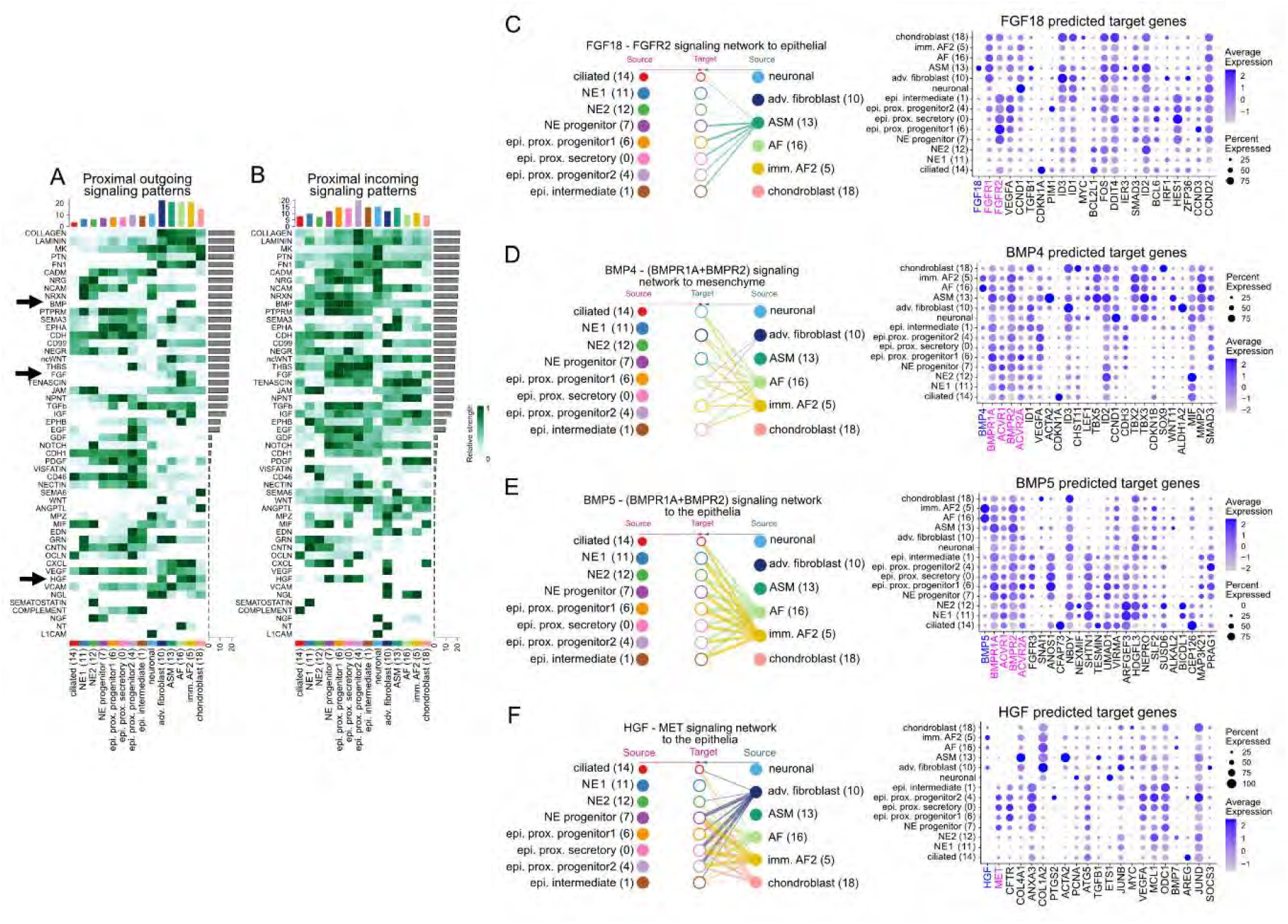
Interactome analyses in the proximal lung neighborhood. **(A-B)** Heatmaps of the significant outgoing (A) and incoming (B) signaling patterns between proximal epithelial identities and their surrounding mesenchyme, as defined in Fig. 1D. The bars represent the outgoing/incoming overall potential on each cluster (top) and pathway (right), predicted by CellChat. **(C)** (left) CellChat hierarchical plot of the FGF18-FGFR2 communication pattern to epithelial clusters. (right) balloon-plot of the top-20 NicheNet predicted target-gene expression levels. **(D-F)** As, as in “C” for (D) BMP4, (E) BMP5 and (F) HGF ligands. Balloon size: percent of positive cells. Color intensity: scaled expression. Ligands: blue, Receptors: magenta.

